# Effect of interhemispheric zero-phase entrainment of the intrinsic mu-rhythm on behavioral and neural markers of predictive coding

**DOI:** 10.1101/2024.05.07.592996

**Authors:** Kirstin-Friederike Heise, Geneviève Albouy, Nina Dolfen, Ronald Peeters, Dante Mantini, Stephan P. Swinnen

## Abstract

Goal-directed behavior requires the integration of information from the outside world and internal (somatosensory) sources about our own actions. Expectations (or ‘internal models’) are generated from prior knowledge and constantly updated based on sensory feedback. This optimized information integration (’predictive coding’) results in a global behavioral advantage of anticipated action in the presence of uncertainty. Our goal was to probe the effect of phase entrainment of the sensorimotor mu-rhythm on visuomotor integration. Participants received transcranial alternating current stimulation over bilateral motor cortices (M1) while performing a visually-guided force adjustment task during functional magnetic resonance imaging. Inter-hemispheric zero-phase entrainment resulted in effector-specific modulation of performance precision and effector-generic minimization of force signal complexity paralleled by BOLD activation changes in bilateral caudate and increased functional connectivity between the right M1 and contralateral putamen, inferior parietal, and medial temporal regions. While effector-specific changes in performance precision were associated with contralateral caudate and hippocampal activation decreases, only the global reduction in force signal complexity was associated with increased functional M1 connectivity with bilateral striatal regions. We propose that zero-phase synchronization represents a neural mode of optimized information integration related to internal model updating within the recursive perception-action continuum associated with predictive coding.

## Introduction

To plan and flexibly adjust goal-directed behavior, information from various external and internal afferent sources must be integrated. Expectations (or ‘internal models’) are generated from prior knowledge and constantly updated based on sensory feedback^1,2^. This ’predictive coding’ implies optimized information integration, resulting in a global behavioral advantage of anticipated action in the presence of uncertainty through minimized variational energy (or complexity)^3,4^.

A fundamental mechanism supporting information integration is the precisely timed functional coupling of neural activity across distant brain areas^5–7^. Specifically coupling with zero-phase shift (i.e., zero-phase synchronization) between remote brain areas is suggested to facilitate neural processing at higher temporal precision^8^, and instantaneous bidirectional^6^ and long-distance interregional neural communication^9,10^. In the sensorimotor network, this information flow between remote homologue motor cortices is reflected by the mutual involvement of bilateral motor areas, even in unimanual behavior^11,12^. Here, our goal was to probe the effect of entraining the intrinsic sensorimotor mu-rhythm (8-13Hz)^13^ bilaterally to zero-phase synchronization between the primary motor cortices on information processing and integration during visuomotor processing^9^. For this purpose, we used transcranial alternating current stimulation (tACS), which has been shown effective in phase and frequency entrainment of intrinsic neural oscillations to the externally applied rhythm^14,15^. As a prototypical example of visuomotor processing, we designed a visually-guided force adjustment task with a constant level of uncertainty (i.e., no sequential regularity), real-time (visual) feedback, and a systematic variation of the effector (uni-/bimanual). According to the ‘predictive coding’ framework, we reasoned that expectations about the upcoming target force based on prior experience and their validation in view of the somatosensory and visual feedback about the actual force applied would lead to subsequent updating of the ‘internal model’. We assumed that a change in predictive coding, i.e., reflecting the level of confidence based on the internal model, would be visible in a global (effector-independent) change in force generation. Conversely, more effector-specific changes in force adjustment would likely represent other than global strategy-related mechanisms, e.g., influenced by the asymmetric distribution of motor skill due to hemispheric dominance. Consequently, we hypothesized that facilitated predictive coding under zero-phase lag stimulation would be reflected by lower uncertainty (or complexity) here quantified with Sample Entropy^16^ of the force signal. We anticipated that this global behavioral change would be paralleled by altered activations across a wider recursive brain network composed of frontal, insular, temporoparietal, striatal, and cerebellar regions^17,18^. On the other hand, if motor-cortical phase entrainment was selectively affecting motor execution, this should be reflected by a more locally confined change in precision (measured with Root Mean Square Error). Finally, we hypothesized that if the two behavioral readouts present the same underlying mechanisms, they would change in the same direction under external phase synchronization. To trace the tACS effect on the behavioral and neural correlates of visuomotor force adjustment, participants received tACS synchronously with task performance during functional magnetic resonance imaging (fMRI) assessing blood oxygen level-dependent (BOLD) signal in the whole brain (overview of experimental procedures shown in Figure 1).

**Figure 1.**
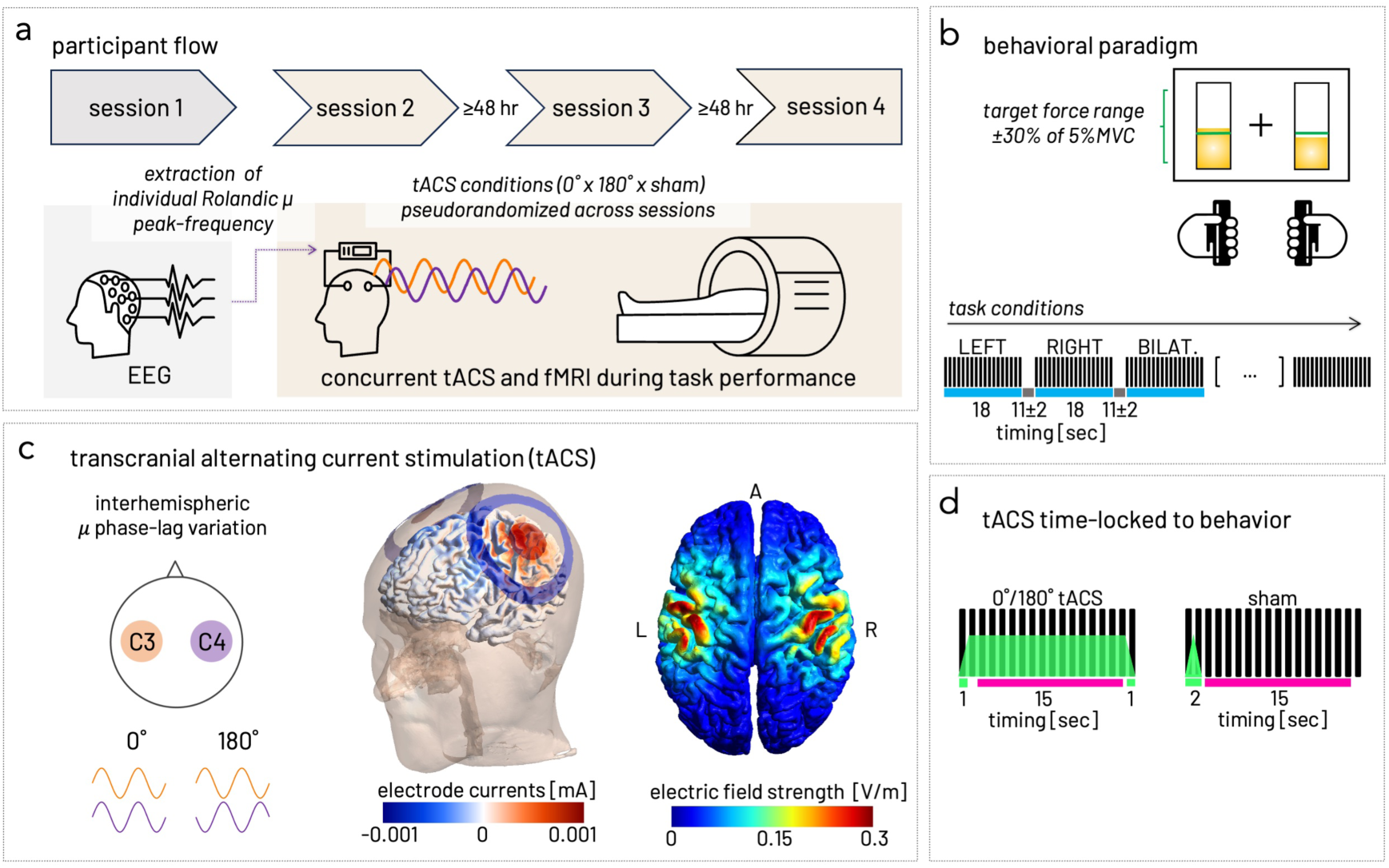
Experimental procedure. **a)** Participant flow through the study. The EEG recording in session 1 served to extract the individual mu peak frequency used as stimulation frequency in the subsequent concurrent tACS and MRI sessions. In a within-subject design, two verum tACS conditions (0°, 180° phase lag) and sham stimulation were pseudorandomized across three sessions. **b)** Visuomotor force adjustment paradigm. Participants were required to adjust their whole-hand isometric grip force to a changing target force level upon a visual cue (green horizontal bar). Using two force transducers, isometric whole-hand grip force was adjusted relative to the participants’ individual maximum voluntary contraction MVC for each hand separately. The target force ranged from 70 to 130% of 5% MVC (green range) in steps of 10% and was presented with a green horizontal bar. Participants received real-time feedback about the actual force exerted (yellow bars). One block consisted of 18 trials of target-force level variations at 1Hz frequency of the same task condition (i.e., LEFT, RIGHT, or BILATERAL, black vertical bars indicate single trials). Task blocks of 18 seconds (blue horizontal bars) were interleaved with rest blocks of varying length (11±2 seconds, grey bars). During the last second of the rest blocks, the task condition cue (arrow to the left, right, or bilateral) was shown. **c)** Interhemispheric mu phase-lag entrainment with bifocal HD-tACS was applied either with 0°- or 180°-degree phase lag between left and right M1 (centered on C3 and C4 positions of the international 10-20-system). Electrical field simulation for an exemplary participant showed focalized fields under the electrodes with maximum intensities between the connector positions of the center and the surrounding ring electrodes. **d)** Verum tACS was controlled against sham stimulation and applied time-locked to the task blocks. During each 18-second task block, stimulation intensity was ramped up and down over one second (green bars). For verum tACS, ramping up/down happened during the first and last trials, resulting in 16 seconds at full intensity (green trapezoid overlay). For sham stimulation, the stimulation was ramped down directly after the 2mA peak-to-peak intensity was reached, resulting in 2 seconds of in- and decreasing stimulation. For the analysis, only data acquired during full electrical current intensity was included (i.e., magenta bars highlight the resulting trials 3-17 extracted from each block for analysis of BOLD signal time series and behavioral data).

Our findings demonstrate that bifocal tACS, entraining the homologous primary motor cortices to zero- (but not anti-) phase lag, leads to a local (effector-specific) modulation of performance precision in contrast to a global (effector-generic) reduction of force signal complexity. Our fMRI results show converging evidence that these distinct behavioral effects are associated with defined BOLD activation changes in bilateral striatal and hippocampal regions. Yet, only the global reduction in complexity of sampled force traces is associated with alteration of functional connectivity between primary motor cortex and striatal and hippocampal regions, implying an induced modulation of information integration within recursive cortico-subcortical loops.

## Results

We used concurrent transcranial alternating current stimulation (tACS) and functional magnetic resonance imaging (fMRI) to probe the effect of phase entrainment of the intrinsic sensorimotor mu-rhythm (8-13Hz)^13^ on the behavioral and neural correlates of visuomotor force adjustment. A bifocal electrode montage was used to apply mu-tACS with 0° phase lag (tACS_0°_), which was controlled against 180° phase lag (tACS_180°_) or sham stimulation in a within-subject cross-over design (three MRI sessions). The stimulation frequency was individualized using the Rolandic mu-peak frequency (range: 9.19 - 12.46 Hz, mean ±SD: 10.65 ± 0.92 Hz, supplemental Figure s1) estimated based on the EEG data collected prior to the MRI sessions (Figure 1a). Complete data from all sessions were collected from 34 participants (19 women, 23.0 ±1.1 years median ± 95% CI), all right-handed (100.0 ±15.3 median ± 95% CI) as evaluated with the Edinburgh Handedness Inventory^19^.

### Interhemispheric zero-phase entrainment induced effector-generic reduction in force signal complexity and effector-specific increase in precision

To identify global (effector-generic) changes in behavioral strategy with altered predictive coding in contrast to effector-specific changes, we characterized behavior (Figure 1b) with two distinct features: 1) Force signal complexity, quantified with Sample Entropy (SmpEn)^16,20^ with smaller SmpEn indicating lower complexity, and 2) Precision, quantified with Root Mean Square Error (RMSE) with higher RMSE indexing larger deviation of the actual force from the target force, i.e., more over- and undershoot (Figure 2a). To investigate the effect of experimentally manipulating the phase lag of mu-tACS between the homologous primary motor cortices on behavior, we used separate linear mixed-effects models (LME, fitted with REML and Satterthwaite approximation) testing the interaction of TASK (left, right, bilateral) and STIMULATION (sham, tACS_180°_, tACS_0°_) conditions on force adjustment RMSE and SmpEn (Figure 2a). This analysis revealed both task- and stimulation-specificity for both behavioral outcomes. Precisely, TASK (F_(2, 46141)_ = 66.95, p<.0001), but not STIMULATION (F_(2, 46064)_ = 2.52, p=.08) impacted force-tracking RMSE. RMSE was expectedly higher in the left and bimanual task conditions compared to the dominant right-hand condition reflecting the influence of hand-dominance on dexterity. The stimulation condition significantly modulated RMSE in a task-specific way (F_(4, 46141)_ = 4.33, p<.005). In the absence of any stimulation effect on bilateral RMSE, this TASK by STIMULATION interaction was driven by a marked reduction in left-hand RMSE (LEFT sham vs. tACS_0°_ ΔEMM = 0.80, -0.275 SE, p_adjusted_ <.001) in addition to an increase in right-hand RMSE (RIGHT sham vs. tACS_0°_ ΔEMM = -0.42, -0.269 SE, p_adjusted_ <.05) under tACS_0°_ (Figure 2b). This stimulation- and effector-specific effect under 0° eliminated the typical performance difference between the non-dominant left and the dominant right hand seen under sham and tACS_180°_ (full results of standardized marginal means contrasts given in Table s2).

**Figure 2.**
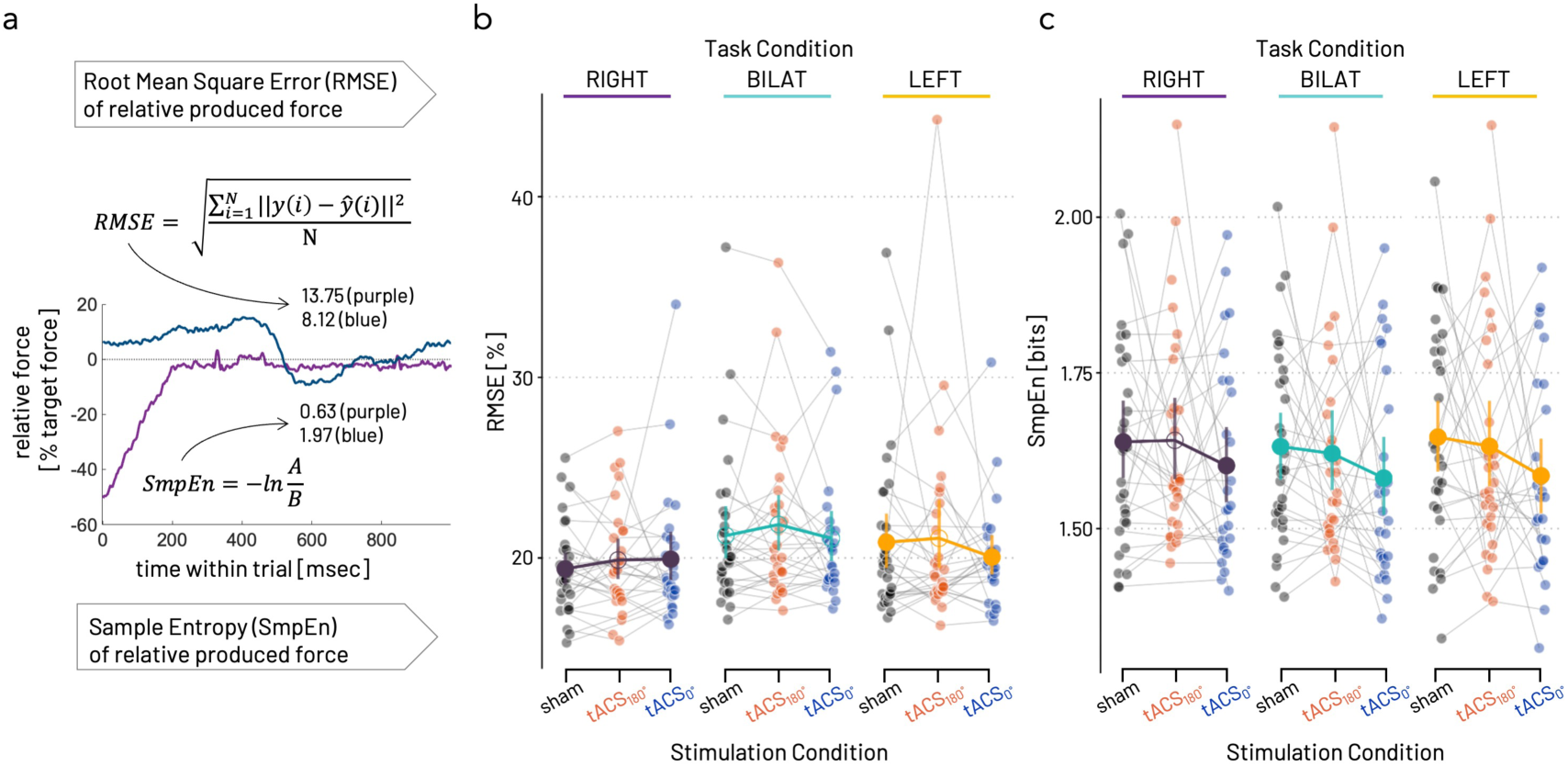
Phase-lag dependent modulation of visuomotor force-tracking behavior with bifocal tACS in the individual mu-peak frequency. **a)** Root mean square error (RMSE) and Sample Entropy (SmpEn) of two example force traces within one trial (1000msec). Sample Entropy(m, r, N) is calculated as the negative natural logarithm of the conditional probability that two vectors (A and B) of m sampling points of a force time series of the length N remain similar at the next point (m+1) within a tolerance of r^20,21^. While the blue force trace has a relatively lower RMSE (8.12%) and higher SmpEn (1.97 bits), the purple trace shows the opposite (13.75% RMSE, 0.63 bits SmpEn). **b)** Modulation of RMSE by stimulation condition (black – sham, red – tACS_180°_, blue – tACS_0°_) within task condition. Individual data points represent averages over trials within participant. Group averages (±95% CI) are overlayed as bigger circles (filled circles indicate significant within-task condition contrasts of stimulation condition). Effector- and stimulation-condition specific modulation of left-hand (yellow) force-tracking RMSE under tACS_0°_, reaching the precision level of the dominant right hand (purple), while right-hand becomes more erroneous under 0° phase lag tACS, in the absence of significant modulation of bilateral (turquoise) RMSE. **c)** Modulation of SmpEn by stimulation condition within task condition, color coding as in b). Stimulation-specific modulation of force tracking SmpEn significantly reduces the complexity of the force time series (i.e., lower SmpEn) under tACS_0°_ independent of the effector. SmpEn under tACS_180°_ is reduced in left and bilateral but not right task conditions.

In the case of force-tracking SmpEn, the LME showed a main effect of TASK (F_(2, 46141)_ = 12.32, p<.0001) with overall higher SmpEn in the right-hand and lower in the bilateral condition, and STIMULATION (F_(2, 46172)_ = 206.08, p<.0001), with lowest SmpEn under tACS_0°_ and highest under sham. As for RMSE, TASK was significantly modulated by STIMULATION (F_(4, 46141)_ = 2.68, p<.05, Figure 2c). However, this interaction was driven by a pronounced reduction of SmpEn under tACS_0°_ for all TASK conditions (sham vs. tACS_0°_, LEFT ΔEMM = 1.644, -0.274 SE, p<001, RIGHT ΔEMM = 0.758, -0.267 SE, p<.001, BILAT ΔEMM = 1.215, -0.267 SE, p<.001). While tACS_180°_ also tended to SmpEn, this effect varied across TASK conditions (sham vs. tACS_180°_, LEFT ΔEMM = -0.21, -0.42 SE, p<.001, RIGHT ΔEMM = -0.88, -0.37 SE, p>.9, BILAT ΔEMM = -0.35, -0.38 SE, p<.001, full results in Table s3).

In summary, tACS_0°_ improved unilateral left-hand force-tracking both in terms of precision, indexed by a reduction in RMSE, as well as in complexity, as indexed by lower SmpEn, whereas tACS_180°_ did not lead to marked behavioral changes within task condition. Importantly, the beneficial effect of tACS_0°_ on RMSE was effector-specific, i.e., the right hand showed the opposite effect (i.e., increase in RMSE), whereas the reduction in SmpEn under tACS_0°_ was effector-independent (i.e., reduced SmpEn for all task conditions), thus potentially indicating two distinct processes. Since the stimulation effects on bilateral behavior largely mirrored the behavioral advantage under tACS_0°_ seen in the unilateral left-hand condition for both outcomes, we focused on the unilateral (left vs. right) contrasts in the subsequent analysis steps.

### Interhemispheric zero-phase entrainment induced an effector-specific modulation of activity in striatal and fronto-parietal cortical regions

To test whether modifying the phase lag of the sensorimotor mu rhythm between homologous primary motor cortices was accompanied by specific neural changes within the regions associated with predictive coding, we next analyzed BOLD signal changes for all possible TASK by STIMULATION condition contrasts. We here report the unilateral (left vs. right) task contrasts, for completeness, the contrasts including the bimanual task condition are detailed in the supplemental material (Tables s4 and s5).

First, we examined which brain regions showed a differential modulation of BOLD signal with tACS_0°_ compared to sham stimulation during left-hand in contrast to right-hand performance. This analysis revealed that neural activity changes for left- and right-hand performance significantly differed in the tACS_0°_ phase lag condition in left and right Caudate from that in the sham condition (Table 1a). Specifically, left-hand performance under tACS_0°_ was associated with lower BOLD activation in the left and right Caudate compared to right-hand performance, whereas the opposite pattern was observed in bilateral Caudate nuclei under sham (Figure 3a).

**Figure 3.**
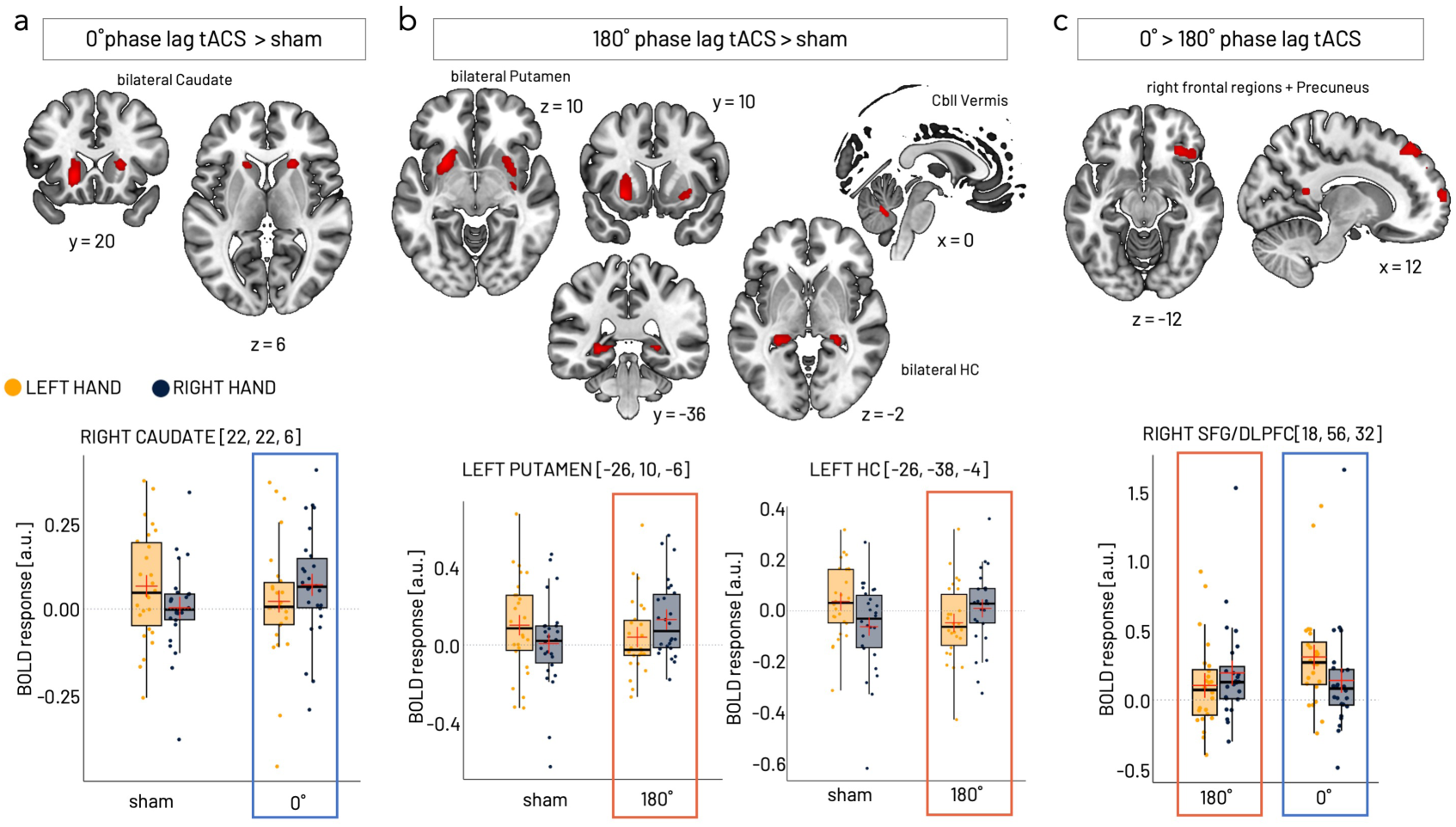
Phase-specific BOLD activation changes during unimanual left versus right-hand force adjustment. **a)** Relative to sham, the activation in bilateral caudate was reduced for left-hand (yellow) in contrast to right-hand (grey/black) force adjustment under tACS_0°_. **b)** Under tACS_180°_, relatively reduced activations during left-hand in contrast to right-hand force modulation were observed for bilateral putamen, bilateral hippocampal, and cerebellar vermis regions. Results are the same when removing the extreme values (Table s6). **c)** Compared to tACS_180°_, left-hand force adjustment was paralleled by relatively higher activation in the right superior frontal gyrus (SFG)/dorsolateral prefrontal cortex (DLPFC) and precuneus under tACS_0°_. Significant clusters of contrast activation maps are shown at a threshold of p_unc-SVC_ < .001 overlaid on the T1-weighted MNI-152 template image included in MRIcroGL (https://github.com/rordenlab/MRIcroGL). Parameter estimates of BOLD signal responses for TASK x tACS interactions (in arbitrary units, a.u.) are displayed as individual data points for within-subject averages. Group statistics shown as boxplots (lower/upper whiskers represent smallest/largest observation greater than or equal to lower hinge ± 1.5 * inter-quartile range (IQR), lower/upper hinge reflects 25% / 75% quantile, lower edge of notch = median − 1.58 * IQR/ sqrt(n), middle of notch reflects group median) and red crosses (group mean). Grey dotted lines demark 0 on the y-axis. Frames highlight verum conditions with tACS_0_, (blue) and tACS_180°_ (orange) consistent with Fig. 1.

**Table 1.**
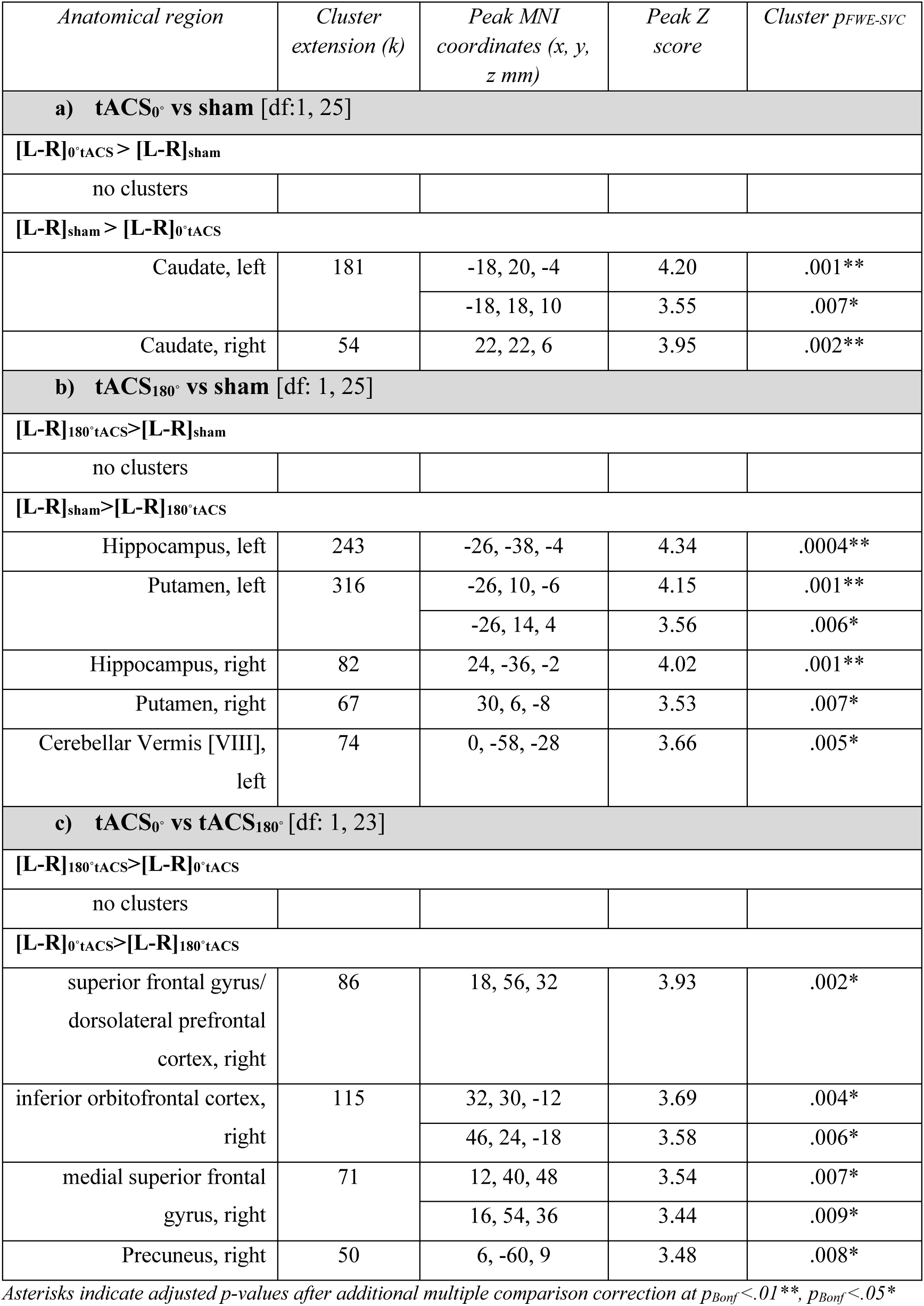
Task- and tACS-specific BOLD signal changes.

| *Anatomical region* | *Cluster extension (k)* | *Peak MNI coordinates (x, y, z mm)* | *Peak Z score* | *Cluster $p_{FWE-SVC}$* |
| --- | --- | --- | --- | --- |
| **a) tACS<sub>0°</sub> vs sham [df:1, 25]** |  |  |  |  |
| **[L-R]<sub>0°tACS</sub> > [L-R]<sub>sham</sub>** |  |  |  |  |
| no clusters |  |  |  |  |
| **[L-R]<sub>sham</sub> > [L-R]<sub>0°tACS</sub>** |  |  |  |  |
| Caudate, left | 181 | -18, 20, -4 | 4.20 | .001** |
|  |  | -18, 18, 10 | 3.55 | .007* |
| Caudate, right | 54 | 22, 22, 6 | 3.95 | .002** |
| **b) tACS<sub>180°</sub> vs sham [df: 1, 25]** |  |  |  |  |
| **[L-R]<sub>180°tACS</sub> > [L-R]<sub>sham</sub>** |  |  |  |  |
| no clusters |  |  |  |  |
| **[L-R]<sub>sham</sub> > [L-R]<sub>180°tACS</sub>** |  |  |  |  |
| Hippocampus, left | 243 | -26, -38, -4 | 4.34 | .0004** |
| Putamen, left | 316 | -26, 10, -6 | 4.15 | .001** |
|  |  | -26, 14, 4 | 3.56 | .006* |
| Hippocampus, right | 82 | 24, -36, -2 | 4.02 | .001** |
| Putamen, right | 67 | 30, 6, -8 | 3.53 | .007* |
| Cerebellar Vermis [VIII], left | 74 | 0, -58, -28 | 3.66 | .005* |
| **c) tACS<sub>0°</sub> vs tACS<sub>180°</sub> [df: 1, 23]** |  |  |  |  |
| **[L-R]<sub>180°tACS</sub> > [L-R]<sub>0°tACS</sub>** |  |  |  |  |
| no clusters |  |  |  |  |
| **[L-R]<sub>0°tACS</sub> > [L-R]<sub>180°tACS</sub>** |  |  |  |  |
| superior frontal gyrus/<br>dorsolateral prefrontal<br>cortex, right | 86 | 18, 56, 32 | 3.93 | .002* |
| inferior orbitofrontal cortex,<br>right | 115 | 32, 30, -12 | 3.69 | .004* |
|  |  | 46, 24, -18 | 3.58 | .006* |
| medial superior frontal<br>gyrus, right | 71 | 12, 40, 48 | 3.54 | .007* |
|  |  | 16, 54, 36 | 3.44 | .009* |
| Precuneus, right | 50 | 6, -60, 9 | 3.48 | .008* |

Asterisks indicate adjusted *p*-values after additional multiple comparison correction at $p_{Bonf} < .01^{**}$ , $p_{Bonf} < .05^{*}$

Next, we tested which brain regions showed a differential modulation of BOLD signal with tACS_180°_ relative to sham stimulation during left-hand in contrast to right-hand performance. In the sham condition, left-hand performance was accompanied by relatively higher BOLD activation in bilateral hippocampus, putamen, and cerebellar Vermis VIII compared to right-hand performance (Table 1b, Figure 3b). This effect changed with tACS_180°_ in all regions. Specifically, bilateral hippocampal activity was significantly reduced for left- as compared to right-hand force adjustment under tACS_180°_ compared to sham. In the left putamen, BOLD activation was relatively lower for the left-compared to right-hand force adjustment. In the right putamen, BOLD activation was increased for right-hand performance under tACS_180°_ compared to the sham condition. Likewise, tACS_180°_, as compared to sham, resulted in a relatively reduced activation in the cerebellar Vermis with left-hand performance as compared to right-hand performance.

Finally, the direct comparison of the two phase-lag variations regarding differential task-specific activations revealed a relative increase in activation for left-in contrast to right-hand performance under tACS_0°_ as compared to tACS_180°_ in right prefrontal, inferior frontal, and superior frontal regions as well right precuneus (Table 1c). Under tACS_180°_ as compared to tACS_0°_, the opposite pattern was observed, i.e., the right-hand force adjustment led to relatively higher activation in these right frontoparietal set of regions (Figure 3c).

Taken together, tACS_0°_ compared to sham stimulation was accompanied by a relative reduction of activation in bilateral caudate during left-hand force adjustment and an increased bilateral caudate activation when the right hand was acting. Under tACS_180°_ as compared to sham, a relatively reduced activation in bilateral putamen, hippocampus, and cerebellar vermis paralleled left-hand force-adjustment, while right-hand performance was accompanied by increased activation in these regions. When compared directly, tACS_0°,_ in contrast to tACS_180°,_ resulted in a relative activation increase during left-hand and a relative activation decrease during right-hand force adjustment in right frontal regions and right precuneus.

### Interhemispheric zero-phase entrainment induced an effector-specific increase of connectivity in striato-motor networks

Further, we were interested in the effect of stimulation on the functional connectivity of the stimulated motor cortical regions with the rest of the brain. Therefore, we performed separate psychophysiological interaction (PPI) analyses^22^ using left and right M1 as seed regions (Figure 4a). These analyses revealed stronger connectivity between the right M1 stimulation target and the contralateral (left) putamen for the left (in contrast to the right) hand performance under tACS_0°_ compared to sham stimulation (Table 2a, Figure 4b). When contrasting tACS_0°_ against tACS_180°_, we also found a relative connectivity increase with the right M1 seed regions under 0° phase lag. Specifically, we found greater connectivity between the right M1 stimulation target and contralateral (i.e., left) putamen, pallidum, inferior parietal, and medio-temporal lobes for left- in contrast to right-hand performance under 0° as compared to tACS_180°_ (Table 2c, Figure 4c). No systematic alteration of task-related connectivity with the left M1 as seed region was found.

**Figure 4.**
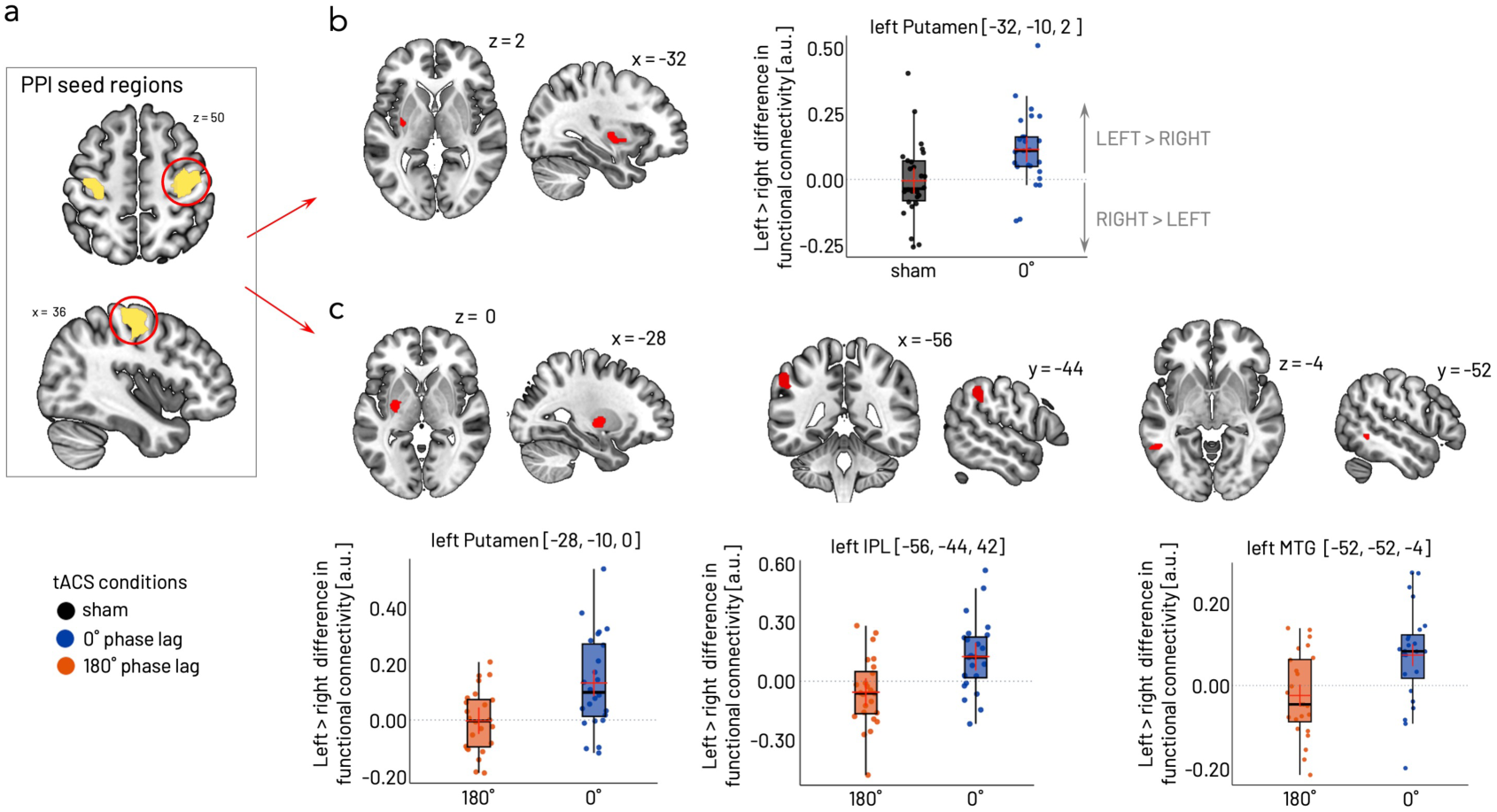
Phase-specific modulation of functional connectivity (PPI) with stimulation target seed in right M1. **a)** Left and right M1 seed regions for PPI analysis (yellow overlays) based on the peak activations on group level resulting from the left-versus right-hand task contrast pooled over all stimulation conditions at p_FWE_ <.05. Only the right M1 seed (encircled in red) yielded significant task-and phase-specific connectivity changes. **b)** Increased functional coupling between right M1 seed and left Putamen under tACS_0°_ (blue) in contrast to sham (black/grey) stimulation was driven by relatively strengthened coupling during left- in contrast to right-hand force adjustment indicated by a positive difference in connectivity. **c)** Increased coupling between right M1 seed and left Putamen (left), left inferior parietal lobule (IPL, middle), and left medial temporal gyrus (MTG, right) during left-hand compared to right-hand task under tACS_0°_ (blue) in contrast to tACS_180°_ (orange). Boxplots and distributions shown in b) and c) depict the relative increase in coupling between indicated areas for the left-hand task compared to the right-hand task. Significant clusters of contrast connectivity maps are shown at a threshold of p_unc-SVC_ < .001 overlaid on the T1-weighted MNI-152 template image included in MRIcroGL (https://github.com/rordenlab/MRIcroGL). Parameter estimates of BOLD signal responses for TASK x tACS interactions (in arbitrary units, a.u.) are displayed as individual data points for within-subject averages. Group statistics shown as boxplots (lower/upper whiskers represent smallest/largest observation greater than or equal to lower hinge ± 1.5 * inter-quartile range (IQR), lower/upper hinge reflects 25% / 75% quantile, lower edge of notch = median − 1.58 * IQR/ sqrt(n), middle of notch reflects group median) and red crosses (group mean). Grey dotted lines demark 0 on the y-axis, i.e., no difference between left- and right-task performance.

**Table 2.** Task-and tACS-related changes in functional connectivity with the right M1 seed region (PPI analysis results)

| *Anatomical region* | *Cluster extension (k)* | *Peak MNI coordinates (x, y, z mm)* | *Peak Z score* | *Cluster p<sub>FWE-SVC</sub>* |
| --- | --- | --- | --- | --- |

| a) tACS<sub>0°</sub> vs sham for left- versus right-hand<sup>§</sup><br>[df: 1, 25] |  |  |  |  |
| --- | --- | --- | --- | --- |
| Putamen, left | 52 | -32, -10, 2 | 3.63 | .005** |
| b) tACS<sub>180°</sub> vs sham for left- versus right-hand<br>[df: 1, 25] |  |  |  |  |
| no clusters |  |  |  |  |
| c) tACS<sub>0°</sub> vs tACS<sub>180°</sub> for left- versus right-hand [df: 1, 23] |  |  |  |  |
| Putamen, left | 132 | -28, -10, 0 | 3.66 | .005* |
|  |  | -22, -4, 4 | 3.32 | .015 |
| Inferior Parietal Lobe, left | 118 | -56, -44, 42 | 3.63 | .006* |
| Medial Temporal Gyrus, left | 32 | -52, -52, -4 | 3.55 | .008* |
| Precuneus, right | 17 | 4, -58, 12 | 3.44 | .009* |

<sup>§</sup> If not shown, the reversed contrast (+T or -T) yielded no significant clusters. Asterisks indicate adjusted p-values after additional multiple comparison correction at $p_{Bonf} < .01^{**}$ , $p_{Bonf} < .05^{*}$

In summary, tACS_0°_ leads to effector-specific (i.e., for left compared to right hand) increased functional connectivity between the stimulation target in right M1 and left hemispheric putamen, inferior parietal, and medial temporal regions relative to the other stimulation conditions.

### Effector-specific behavioral changes are linked to distinct cortical and subcortical BOLD activation changes contrasting effector-generic behavioral changes associated with increased functional cortico-striatal connectivity

To further disentangle potentially different mechanisms underlying the distinct effects of tACS_0°_ visible in the two behavioral outcome parameters with an effector-specific variation in RMSE (i.e., reduction in left- and increase in right-hand RMSE) and an effector-generic reduction in SmpEn, we ran a set of separate second-level regression models. For this purpose, stimulation condition differences were computed for each hand separately for the two behavioral outcome parameters (i.e., ΔRMSE = RMSE_0°_ - RMSE_sham_, ΔSmpEn = SmpEn_0°_ - SmpEn_sham_) and used as covariates to test the brain-behavior relationships with the BOLD activation and functional connectivity (PPI).

We found that a stronger reduction of left-hand (but not right-hand) RMSE was associated with a greater reduction in right caudate and right hippocampal activation under tACS_0°_ as compared to the sham condition (Table 3a, Figure 5a). In contrast, the increase in right-hand RMSE was associated with stronger activation in the right posterior cingulate under tACS_0°_ compared to sham (Table 3b, Figure s7). The relative reduction of left-hand SmpEn under tACS_0°_ as compared to the sham condition was significantly associated with increased functional connectivity between the right M1 seed and right caudate, left putamen, and to a lesser extent, with the right cerebellar Crus I (Table 3g, Figure 5b). For the right hand, lower SmpEn was likewise associated with a connectivity increase between the right M1 and right putamen (Table 3h).

**Figure 5.**
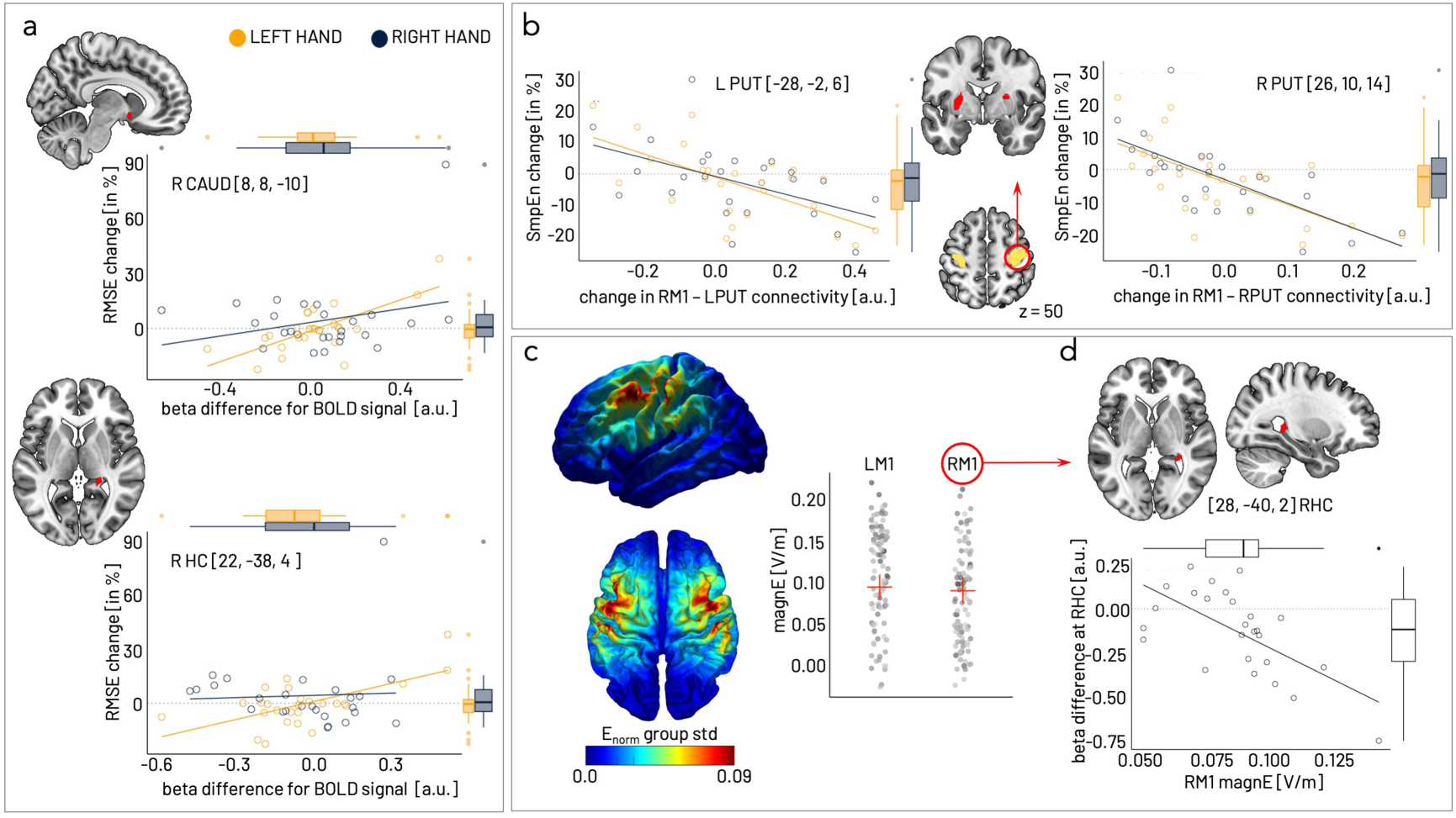
Associations between neural and behavioral changes under 0° phase lag tACS. **a)** Reduced left-hand (yellow) but not right-hand (grey) RMSE under tACS_0°_ was associated with a relative decrease of right caudate (top) and right hippocampus (bottom) activation. Association between right-hand RMSE increase and right PPC activation increase is shown in Figure s7. **b)** The reduction in SmpEn under tACS_0°_ was associated with increased connectivity between right M1 seed region and bilateral striatal regions, here shown for left and right putamen. This regression was significant for both left- and right-hand SmpEn. **c)** Group average of the electrical field (E_norm_). Individual variation of electrical field strength at left and right M1 seed, i.e., stimulation targets. **d)** Association between higher eField strength in RM1 and more pronounced reduction in functional RM1-RHC connectivity (negative difference of beta values tACS_0°_ - beta values sham, Figure s8) for left-versus right-hand contrast). Significant clusters of contrast connectivity shown at a threshold of p_unc-SVC_ < .001 overlaid on the T1-weighted MNI-152 template image included in MRIcroGL (https://github.com/rordenlab/MRIcroGL). R CAUD right caudate, R HC right hippocampus, L/R PUT left/right putamen, L/R M1 left/right primary motor cortex.

**Table 3.** Regression analyses of associations for tACS_0°_ versus sham contrasts.

| *Anatomical region* | *Cluster extension (k)* | *Peak MNI coordinates (x, y, z mm)* | *Peak Z score* | *Cluster $p_{FWE-SVC}$* |
| --- | --- | --- | --- | --- |
| **a) $BOLD [L tACS_0^+ > L sham] \times RMSE [R tACS_0^+ > R sham]^S$** |  |  |  |  |
| Caudate, right | 17 | 8, 8, -10 | 3.69 | .004** |
| Hippocampus, right | 8 | 22, -38, 4 | 3.53 | .007* |
| **b) $BOLD [R tACS_0^+ > R sham] \times RMSE [R tACS_0^+ > R sham]^S$** |  |  |  |  |
| Posterior Cingulate, right | 27 | 12, -42, 18 | 3.56 | .006** |
| **c) $BOLD [L tACS_0^+ > L sham] \times SmpEn [L tACS_0^+ > L sham]^S$** |  |  |  |  |
| no clusters |  |  |  |  |
| **d) $BOLD [R tACS_0^+ > R sham] \times SmpEn [R tACS_0^+ > R sham]^S$** |  |  |  |  |
| no clusters |  |  |  |  |
| **e) $PPI (RM1 seed) [L tACS_0^+ > L sham] \times RMSE [R tACS_0^+ > R sham]^S$** |  |  |  |  |
| no clusters |  |  |  |  |
| **f) $PPI (RM1 seed) [R tACS_0^+ > R sham] \times RMSE [R tACS_0^+ > R sham]^S$** |  |  |  |  |
| no clusters |  |  |  |  |
| **g) $PPI (RM1 seed) [L sham > L tACS_0^+] \times [L sham > L tACS_0^+]^S$** |  |  |  |  |
| Caudate, right | 52 | 22, 20, 8 | 4.10 | .001** |
| Putamen, left | 46 | -28, -2, 6 | 3.53 | .007* |
| Cerebellum Crus I, right | 4 | 42, -58, -32 | 3.22 | .018 |
| **h) $PPI (RM1 seed) [R sham > R tACS_0^+] \times [R sham > R tACS_0^+]^S$** |  |  |  |  |
| Putamen, right | 77 | 26, 10, 14 | 3.98 | .002** |
| **i) $RM1 eField as covar - (+T) (LvsR 0^\circ vs sham)^S$** |  |  |  |  |
| no clusters |  |  |  |  |
| **j) $PPI (RM1 seed) - RM1 eField as covar - (+T) (LvsR 0^\circ vs sham)^S$** |  |  |  |  |
| Hippocampus, right | 12 | 28, -40, -2 | 3.68 | .005** |

Df: [1, 24] for all contrasts shown in this table. <sup>S</sup> If not shown, the reversed contrast (+T or -T) yielded no significant clusters. Asterisks indicate adjusted *p*-values after additional multiple comparison correction at $p_{Bonf} < .01^{**}$ , $p_{Bonf} < .05^*$

In the last analysis step, we investigated whether the individual variation of the electrical field strength induced by the bifocal electrode montage (Figure 5c) was associated with the stimulation-induced BOLD activation or functional connectivity changes under tACS_0°_ compared to sham. We found no association between BOLD activation changes and the electrical field strength in the left or right M1. However, the electrical field in the right M1 was significantly linked to the stimulation-dependent modulation of functional connectivity between the right M1 seed region and the right hippocampus (Table 3j). Specifically, a stronger electrical field in the right M1 was associated with a more pronounced reduction in functional connectivity between the right M1 and right hippocampus under tACS_0°_ compared to sham as indexed by negative difference values (Figure 5d, Figure s8). For completeness, we also tested the associations with the electrical field strength in the left M1, which revealed the same trend, i.e., higher electrical field strength led to a more pronounced reduction in coupling between the left stimulation target and hippocampal and striatal regions under tACS_0°_ compared to sham (Table s9, Figure s10).

In summary, the reduction in RMSE seen in the left hand under tACS_0°_ compared to sham was accompanied by a relative decrease in right caudate and right hippocampal BOLD activation. In contrast, the relative increase in right-hand RMSE seen under tACS_0°_ was related to an increase in posterior cingulate BOLD activation. The effector-independent reduction in SmpEn was associated with increased functional connectivity between right M1 (seed region) and bilateral striatal regions. In addition, we found an association between higher individual electrical field strength within the right M1 and the magnitude of the relative reduction of functional connectivity between the right M1 and hippocampus under tACS_0°_.

## Discussion

We here asked whether 0°phase lag oscillatory coupling between homologous primary motor (M1) cortices would promote predictive coding, as indexed by reduced variational energy (i.e., complexity) of the behavioral read-out, and if this would be paralleled by changes in brain activation within the network of brain regions associated with predictive coding^17,18^. We assumed that a change in predictive coding, i.e., reflecting the level of confidence based on the internal model, would be visible in a global (effector-independent) change in force generation. Conversely, effector-specific behavioral change would likely reflect other underlying control mechanisms. To test this hypothesis, we used a visually-controlled force adjustment paradigm and characterized the force adjustment behavior with two distinct features, namely the complexity of sampled force signal (SmpEn, Figure 2a) and the quantification of the relative force over- and undershoot (RMSE). While a high level of confidence based on the internal model may serve better performance precision, the two outcome parameters are not intrinsically correlated and thus allow the distinction between systems-level strategy (SmpEn) and effector-specific skill (RMSE). To probe the effect of specificity of phase-locking of interhemispheric motor-cortical oscillatory brain activity, we used a within-subject design and tested tACS_0°_, applied within the individualized sensorimotor mu (8-13Hz) band, against two control conditions (i.e., 180° phase lag and sham) concurrently with task-based fMRI.

We demonstrate that bifocal tACS with zero phase lag (but not tACS_180°_) between homologous motor areas leads to a global (i.e., effector-generic) reduction of force signal complexity, suggestive of facilitated predictive coding which is contrasted against an effector-specific modulation of precision error. These two distinct tACS_0°_-induced behavioral effects are associated with different neural responses. Specifically, as additionally confirmed by regression analyses, effector-specific force precision changes under tACS_0°_ are associated with defined BOLD activation changes in bilateral striatal and hippocampal regions. In contrast, the global reduction in complexity of sampled force traces is associated with altered functional connectivity between the right primary motor cortex seed and bilateral striatal as well as hippocampal regions.

In the behavioral paradigm employed here, the expectations (or internal model) were actively generated during task experience in the first session before the MRI sessions (three runs of 30 blocks with 19 trials of force adjustments each, Figure 1a). Thus, despite a constant level of uncertainty (i.e., no sequential regularity of force levels), a priori knowledge included a defined set of rules, specifically the number of force levels in a narrowly circumscribed force range as well as the online visual feedback about actual force relative to target force allowing for updating the internal model. A strategic change in the integration of available information, e.g., down-weighting of visual feedback information with increasing confidence of the internal model^23^, should translate into a global, less effector-specific, state change, as reflected here by an overall minimized signal complexity of the behavioral read-out^24,25^. On the other hand, improved motor skill would likely be observable in a more selective quantitative change of the behavioral read-out itself, such as seen here in increased precision. Our results support these hypotheses in that the non-dominant left hand showed improved performance precision (i.e., reduced RMSE) most likely because there is generally more room for improvement on the non-dominant, less dexterous side, which outlasted the extensive familiarization session. In contrast, the reduced signal complexity indexed by a decrease in SmpEn was a more global observation visible across all task conditions (i.e., effector-independent) hinting at a strategic change. These results suggest that the two parameters reflect different underlying processes because, otherwise, both should have changed in the same direction with the experimental manipulation of stimulation-induced phase lag.

In parallel with the behavioral effects under tACS_0°_, we observed a phase-lag-specific tACS modulation of task-related brain activity in bilateral striatum as well as right frontal and parietal regions. Specifically, in the tACS_0°_ condition compared to sham, the contrast of left-versus right-hand performance revealed a decrease in BOLD activation in bilateral caudate nuclei. At the same time, increased functional coupling between the right M1 stimulation target and the contralateral (left) anterior part of the putamen was observed during left-relative to the right-hand force adjustment. Although our behavioral paradigm represents a well-trained, highly repetitive task without sequential regularity to be learned, tACS_0°_ induced modulations of activity and connectivity primarily in anterior striatal regions, including the anterior portion of the caudate nucleus, associated with integrating probabilistic regularities^26^. Prefrontal and ventral premotor regions primarily project to the caudate nucleus and the anterior putamen^27^. Reduced caudate activation during left-hand performance under tACS_0°_ may therefore reflect reduced processing requirements of input from prefrontal and premotor regions for left-hand action, which may go along with reduced visuomotor integration load within the ventral premotor cortex going along with an improved internal model. Thus, lower cognitive control may be required due to fewer breaches of expectations, i.e., less surprise due to deviations between anticipated and required force, for the left hand with tACS_0°_ compared to sham^28,29^, and hence lower caudate activation is needed.

Given the projections from the anterior cingulate and prefrontal cortex^27^, the putamen has been suggested as the mediating hub for signals of ‘surprise’ arising from the mismatch between top-down prediction and bottom-up ascending (perceptual) representations^29^. While the anterior striatum, specifically the putamen activation, has been reported to scale with grip force precision in healthy volunteers even without visual feedback^30^, the (particularly left) putamen’s role in prediction error processing based on confidence independent of perceptual feedback has been suggested^31^. Thus, the increased coupling between the right M1 and the left putamen observed under tACS_0°_ for left-compared to right-hand action may indicate a fundamentally changed processing, i.e., re-weighting or attenuation of perceptual feedback under tACS_0°_ relative to sham with an improved internal model. This interhemispheric functional coupling is structurally supported by putative crossing projections from the primary, supplementary, and pre-motor cortical regions onto the contralateral putamen^32,33^ and may add to the global behavioral effect of reduced complexity (i.e., SmpEn) of the sampled force traces under bilateral motor-cortical 0°phase synchronization. Importantly, this hypothesis receives converging support from our finding of an increased association between increased corticostriatal coupling and a more pronounced reduction of SmpEn for either hand.

These connectivity results and the fact that BOLD activation changes were exclusively associated with changes in performance precision (RMSE) substantiate the hypothesis that the behavioral results rely on distinct underlying neural mechanisms. Precisely, a more pronounced reduction in right caudate and hippocampal activation paralleled reduced left-hand RMSE in contrast to stronger increase in right PCC that paralleled the increase in right-hand RMSE. These distinctive associations between RMSE and BOLD signal change are likely to explain the selective improvement of the non-dominant hand precision, going along with reduced caudate and hippocampal processing requirements. In contrast, a slightly worsened precision of the dominant hand may be associated with an increased computation of expectations in posterior cingulate cortex^34^, as suggested by our regression results. Our findings of a significant association between higher electrical field strength within the right motor-cortical stimulation target and more pronounced modulation of the functional coupling between right M1 and hippocampal regions, but not directly with BOLD activation, further point to an indirect stimulation effect, i.e., the modulation of network dynamics accessed through the interhemispheric motor-cortical phase synchronization. Specifically, while connectivity increases under tACS_0°_ between the right stimulation target in M1 and left hemispheric putamen, IPL, and MTL, higher electrical field strength in the right stimulation target was associated with reduced intrahemispheric connectivity between the right M1 and hippocampus. Together, these results might indicate that the interhemispheric synchronization and potentially the enslaving of the non-dominant hemisphere to the same rhythms as the dominant one is more relevant for the observed global behavioral change. The reduction of right hemispheric M1-hippocampal connectivity, on the other hand, might reflect the reduction of the search for a sequential pattern with increased confidence in the internal model. The hippocampus potentially functions as a prime computational unit to manage prediction errors thanks to its functional architecture, allowing for the matching of predictive versus erroneous (‘mismatch’) signals within the hippocampal–neocortical loop^35,36^. On the other hand, the lack of a behavioral effect together with the relatively reduced activations in bilateral putamen and hippocampal regions and no effect on functional connectivity under tACS_180°_ compared to sham may indicate less effective integration of top-down predictions with bottom-up perceptual feedback and updating of the internal model under tACS_180°_.

In the direct contrast of both verum conditions, we found relatively higher activity under tACS_0°_ compared to tACS_180°_ for the left-hand relative to right-hand task condition in frontal and parietal regions, hinting at a phase-specific modulation of the frontoparietal network associated with the flexible allocation of attention, task planning, and generation of inferences^38^. Specifically, the right inferior orbitofrontal cortex is associated with the computation of inferences (e.g., rule encoding) and prediction error by integrating information from multiple sensory and association sources (e.g., sensory, visceral, hippocampal, amygdala, striatal) and by exerting descending control over action through its connections with the prefrontal cortex^39,40^. Likewise, relatively higher activation in the right superior frontal gyrus, dorsolateral prefrontal, and dorsomedial prefrontal cortex alongside increased activation in the precuneus point to a more pronounced frontoparietal control under tACS_0°_ (in contrast to tACS_180°_) which is behaviorally beneficial. In support of this notion, the altered weighting of frontal over parietal regions has been found to occur with increased visuomotor experience and is interpreted as being characteristic of more flexible shifting of visuomotor attention^41^. A probable explanation is that a stronger engagement of regions within the frontoparietal control network facilitates the stronger integration of the stimulation targets (i.e., primary motor cortices) within a cortico-striatal loop under tACS_0°_ by generally allowing for more flexible integration of expectation and re-evaluation of perceptual information^42^. At the same time, this state of higher neural coupling may result in a more flexible allocation of attentional control and hence more precise performance (i.e., lower prediction error) for the non-dominant left hand. Furthermore, our results show relatively increased functional coupling between the right M1 stimulation target and contralateral, i.e., left-hemispheric putamen, inferior parietal lobule, and medial temporal gyrus under tACS_0°_ compared to tACS_180°_ for left-compared to right-hand force adjustment. These changes in functional connectivity additionally support a phase-specific alteration in cortico-cortical and cortico-subcortical network organization, with stronger coupling during non-dominant hand action.

In conclusion, we demonstrate distinct phase-specific behavioral effects in visuomotor force adjustment using bifocal tACS to entrain the intrinsic mu rhythm between homologous sensorimotor cortices with 0° phase lag. Specifically, we observed an effector-specific modulation of performance precision and an effector-generic minimization of force signal complexity. These phase-specific effects on behavior were paralleled by effector-specific BOLD activation changes in bilateral caudate as well as increased functional connectivity between the right motor-cortical stimulation target and contralateral putamen, inferior parietal, and medial temporal regions. As confirmed by regression analysis, effector-specific changes in performance precision were associated with contralateral caudate and hippocampal activation decreases, while effector-independent reduction in force signal complexity was associated with increased functional connectivity between the right stimulation target and bilateral striatal regions. Therefore, we propose that the interhemispheric 0° phase lag synchronization likely results in alterations of functional connectivity within a broader network, particularly the cortico-striatal loop, potentially promoting key processes related to internal model updating and re-weighting of feedback information within the recursive perception-action continuum associated with predictive coding. Altogether our findings suggest that focal interhemispheric synchronization can induce activity and connectivity alterations within a broader cortico-subcortical network with distinct effects on behavior.

## Methods

### Participants

Young, healthy participants (N = 38, 19 – 30 years age range) were recruited through flyers and adverts on social media. All participants gave written informed consent and were systematically screened with a structured interview for neurological and psychiatric diseases and contra-indications against non-invasive brain stimulation, MRI scanning, or any other procedure and product applied in this study, conforming with current safety guidelines. The medical ethics committee of KU Leuven approved the protocol and all procedures of this study following the Declaration of Helsinki in its 2008 version (national registration protocol number B322201526159, local protocol number s58333). Participants received reimbursement of 15.-€ per hour.

Three participants dropped out from the study, i.e., one due to an illness unrelated to the study procedures, one did not tolerate the discomfort of the tACS stimulation, and one developed a headache several hours after the first MRI session. Data from a fourth participant was excluded due to technical problems with the task set-up, corrupting data collection.

### Experimental procedures

The participants underwent four consecutive experimental sessions, i.e., one EEG session and three MRI sessions with concurrent tACS (figure 1a). The EEG session (session 1) served to familiarize participants with the behavioral paradigm and to extract their individual mu-peak frequency, which was used as tACS stimulation frequency in the subsequent MRI sessions (sessions 2-4). Across the three MRI sessions, the three stimulation conditions (verum tACS with 0° or 180° phase lag or sham tACS) were applied in a pseudo-randomized order, counter-balanced across participants. To minimize potential carry-over effects between tACS conditions^43,44^, the MRI sessions were separated by 48 hours at minimum.

### Behavioral Paradigm

We used a visuomotor force-tracking task (Figure 1b) which, according to the ‘predictive coding’ framework, translates into expectations about the upcoming target force based on prior experience and their validation given the available somatosensory and visual feedback corresponding to the actual force applied and subsequent updating of the ‘internal model’. Upon a visual cue (a green horizontal line) indicating the target force, the participants had to adjust their grip force to changes in target force level at a rate of 1Hz, either with the left, the right, or both hands simultaneously. The bilateral task condition involved a symmetric adjustment of both hands, i.e., left and right target force cues indicated the same level. The participants’ actual force level was fed back in ‘real-time’ (i.e., with a 20 msec delay) using a yellow bar. Two MR-compatible dynamometers (SS25LB hand dynamometer and the MP160, Biopac Systems Inc., Goleta, CA, USA) were used to perform whole hand isometric grip force. Directly before the fMRI sequence, the grip force was adjusted to 5% of the individual maximum voluntary contraction (MVC) for each hand separately. Target force levels were adjusted with respect to the 5% MVC force level (corresponding to a target force of 100%) and varied randomly in steps of 10% from 70% to 130% of the target force. During one fMRI session, 36 blocks (12 per task condition) of 18 individual force adjustment trials (each of 1-second lengths) were performed. The participants were instructed to follow the target force with their own force adjustment as fast and as precisely as possible and to hold the force constant until the new target force was visible.

Thus, while the range of target force levels was within fixed boundaries, the sequence of target force levels was pseudorandomized and did not repeat the same order of force increments within one experimental session. One block consisted of either left, right, or bimanual force adjustments only. Each task condition was indicated by an arrow (task condition cue) at the start of each practice block and repeated four times, i.e., four blocks of 18 seconds were alternated within each session with fixed order for each participant, which was counterbalanced across sessions. Rest blocks were offered between practice blocks and required the participants to fixate a cross in the middle of the screen for approximately 11 seconds (±2 seconds jitter) without moving or imposing force on the dynamometers. Visual stimuli were programmed in LabVIEW 2018 (National Instruments, Austin, TX, USA), which was also used for logging the performance throughout the experiment for offline analysis. All participants practiced the task in the EEG session preceding the first MR session during three runs of 30 blocks, each consisting of 19 single trials with likewise randomly varying increments of target force.

### Operationalization of performance and behavioral data analysis

Force time series were sampled at 1000Hz and smoothed, offline, using a Savitzky-Golay filter (MATLAB’s ‘sgolay’ function using 5^th^ order, i.e., over a window of 25 samples window). From these force time series, two outcome measures were extracted for each individual trial to describe distinct features of force adjustment (Figure 2a): 1) Root Mean Square Error (RMSE) to characterize the deviation of the actual force from the target force, i.e., over- and undershoot, and 2) sample entropy (SmpEn)^16,20^ to quantify the randomness or uncertainty (in the following referred to as ‘complexity’) of the sampled force time series. Root mean square error (RMSE) was calculated with the MATLAB function ‘rmse’. In the context of Information Theory, Sample Entropy is used to describe the regularity of dynamic processes like biological time series^16^. Sample Entropy(*m, r, N*) is calculated as the negative natural logarithm of the conditional probability that two vectors of *m* sampling points remain similar at the next point (m+1) within a tolerance of a radius threshold *r* largely independent of the overall length of the time series of *N* sampling points^20,21^. The parameters m, r, N are set a priori. Here, we used embedding dimensions m=2 for a time delay of 1 sampling point, with a threshold r=0.2*SD[time series]. Hence, Sample Entropy is a function of the probability of the occurrence of each value of the time series, which translates into low information or complexity (i.e., low SmpEn, Figure 2a purple example force trace) in the case of little change over the time series and high complexity (i.e., high SmpEn, Figure 2a blue example force trace) in case of abrupt changes. Increases in force signal complexity, indexing complete independence of each data point, would result in values near the maximum defined by the base of the natural logarithm, i.e., 2.72. For the bimanual task condition, outcome values (RMSE and SmpEn, respectively) were averaged over the left and right hand for each trial after confirming that stimulation effects on each hand were comparable in direction in the uni- and bilateral task conditions (Figure s11). Separate linear mixed effects (LME) models were fitted to explain the variance in (1) RMSE and (2) SmpEn with TASK (left, right, bilateral), STIMULATION CONDITION (sham, 0°, 180°), and their interaction modeled as fixed effects. Random intercepts were modeled on the participant level (restricted maximum likelihood estimation, REML). Standardized parameter estimates were obtained by fitting the model on a standardized dataset version relative to reference categories (TASK = right, STIMULATION CONDITION = sham). Results for LME models are reported with Type II F-statistics using Satterthwaite approximation for degrees of freedom. Pairwise contrasts of individual factor levels are directly estimated from the LME model and reported as standardized differences (Δ_EMM_, SE) and p-values with Bonferroni-Hochberg adjustment for multiple comparisons^45^. All LME were fitted in R language and environment for statistical computing [https://www.R-project.org/], using the lme4^46^ library together with the ‘easystats’ framework^47^ for reporting (full list of libraries in supplemental code accompanying this paper).

### Transcranial alternating current stimulation (tACS)

Bifocal transcranial alternating current stimulation (tACS) was applied at the individual mu-peak frequency to homologous primary motor cortices (Figure 1c). Two battery-driven MR-compatible electrical stimulators (DC-stimulator Plus with MR extension, neuroConn GmbH, Ilmenau, Germany) were used to apply bifocal tACS. Stimulation on- and off-set was externally triggered by custom software (programmed in LabVIEW 2018 (National Instruments, Austin, TX, USA), time-locked to the behavioral task, and synchronized with the fMRI sequence. The stimulation intensity of ±1mA (i.e., 2mA range) peak-to-peak was applied in trains of 18 seconds, including a ramp-in and ramp-down phase of 1 second. Sham stimulation consisted of the ramp-in and ramp-out phase only and a single cycle at maximum intensity, i.e., it had a total length of approximately 2 seconds.

### Individual mu peak frequency estimation

In the EEG session preceding the MRI sessions, a task-free recording of six minutes (eyes open, attending a fixation cross) was used to estimate the individual Rolandic mu peak frequency (iµPF). Continuous EEG data were acquired with a sampling frequency of 1000Hz, 280Hz cut-off frequency (127 cephalic active surface electrodes referenced to FCz, actiCAP, actiChamp, BrainVision Recorder version 1.21.0004, BrainProducts GmbH, Gilching, Germany). Offline data preprocessing and analysis were performed using EEGLAB (version 2019.1) and Fieldtrip (version 20190419) toolboxes, in addition to the polyfit function implemented in Matlab 2018b. Preprocessing included down-sampling to 250Hz, high-pass filtering at 1Hz using a 1^st^ order Hamming windowed sinc FIR filter, removing line noise (50 and 100 Hz)^48^, automatic eye movement artifact removal using canonical correlation analysis^49^. Fast Fourier transformation with 1 Hanning taper was used on non-overlapping epochs of 2 seconds length to calculate the frequency spectra between 4 and 45Hz. The individual mu peak frequency was estimated using a Gaussian curve fit^50^. Specifically, the highest local maximum within the mu frequency range ±1Hz (i.e., 7 – 14Hz) was estimated with 0.01 Hz resolution for both C3 and C4 electrodes supposed to capture signals from left and right primary motor cortices. Based on visual inspection, iµPF from the hemisphere that showed the most optimal fit was chosen as stimulation frequency (N = 35 from C3, N = 3 from C4). In the final sample, the iµPF ranged from 9.19 to 12.46 Hz (10.65 ± 0.92 Hz, Figure s1).

### Electrode montage for tACS application

A bifocal center-ring high-definition electrode montage (Figure 1c) was used with center electrodes positioned over the left and right primary motor cortex, identified by the electrode position of C3/C4 of the international 10–20 EEG system, each with a surface area of 9 cm^2^, diameter 3.4 cm. The outer ring-shaped patch electrodes were placed with a 2.05 cm distance between electrode borders (individual surface area 35 cm2, inner diameter 7.5 cm, outer diameter 10 cm).

To ensure optimal contact between stimulation electrodes and skin, the skin was prepared with an abrasive paste (Nuprep, Weaver ad company, Aurora, Colorado, USA). Thereafter, electrodes were prepared with a 2mm layer of MR-compatible conductive paste (TEN20, Weaver ad company, Aurora, Colorado, USA and then firmly attached to the scalp and fixated with a standard non-adhesive textile elastic bandage to prevent dislocation throughout the experiment.

MR-compatible cables connected each electrode pair of one hemisphere to a separate tACS stimulator. Phase differences (0° versus 180°) between center electrodes were experimentally manipulated through the phase-offset settings within each stimulator.

### Impedance

Static impedance was checked after electrode placement to establish impedance values <5kOhm for left and right hemispheric electrode pairs independently of each other outside the MRI. Due to the resistors of the MR tACS equipment, dynamic impedance measurements from inside the MRI environment increased by 10kOhm. When the participants were placed inside the MRI bore, the stimulators were connected to the tES stimulator cable extensions in the control room, and the impedance was checked again. During the fMRI sequence with concurrent tACS, the impedance was monitored constantly by one researcher not blinded to the stimulation condition (KFH), and impedance minima and maxima were documented per stimulation train throughout the task-related fMRI sequence.

Given the features of the impedance values (i.e., non-gaussian distribution, bounded range) and the difference in values outside and inside the MRI environment, two separate Bayesian general linear models (Gamma family, identity link) were fitted for measurements taken (1) outside the scanner after electrode placement and (2) inside the scanner before and during the scans. As we wanted to control whether impedance changed over time and whether this was potentially modulated differently by stimulation conditions, we also ran the ‘inside’ model, including the consecutive number of impedance checks as an indicator for progressing time. We found no change in impedance over the time course of the experiment, neither overall nor for the individual stimulation conditions (Tables s12-s14).

### Evaluation of neurosensory side effects of tACS and stimulation condition estimation

Subjective estimation of neurosensory side effects was evaluated with a standardized questionnaire^51,52^. Side effects such as tingling, burning sensation, or pain were rated on a categorical scale ranging from ‘none’ to ‘strong’ after each of the three tACS-MRI sessions. After the last session, participants were asked to give their estimation about whether they received sham or verum stimulation or whether they were unable to differentiate conditions for each of the preceding sessions. Bayesian probability tests were performed to compare the relative frequencies of (1) the reported intensities of the neurosensory side effects for the separate stimulation conditions and (2) the estimation of sham versus verum stimulation conditions. Overall, the volunteers evaluated the intensity of stimulation-related neurosensory side effects as being very low, and this was not different among the three stimulation conditions (Figure s15, Table s16). Based on the stimulation estimation after the last session, we found no evidence to support a difference in the participants’ evaluation of any of the stimulation conditions as being real stimulation. However, participants were more likely to categorize sham and tACS_0°_ as being sham than tACS_180°_ (Table s17, Figure s18). Together these results render neural and behavioral effects less likely to be caused by differences in neurosensory side effects or the participants’ expectations regarding varying effects elicited by the different stimulation conditions.

### Electrical field simulation

For each participant, electrical fields for the bifocal center-surround montage (Figure 1c) were simulated with SimNIBS (v4.0.0)^53^ based on individual finite element head models derived from individual T1-weighted MRI data of the same participant.^54^ Circular (34 mm diameter) and ring electrodes (75/100 mm inner/outer radius) were centered at the C3 and C4 positions of the international 10-20 system registered to native space. All electrodes were modeled as 1.5 mm thick rubber layers (conductivity 0.1 S/m) with 1.5 mm thick layers of conductive gel underneath (conductivity 8 S/m, estimated based on sodium concentration in the TEN20 paste, as stated by the manufacturer). Cable connector positions were explicitly modeled^55^ to reflect their standardized placement in the direction of the 10-20 electrode position Pz for the center and Fz for the outer ring electrodes to yield approximately a 90° angle between connectors. Group average field strengths were computed over all participants. Additionally, local field strengths were extracted for regions of interest for participants of the final sample included in the fMRI data analysis and subjected as covariates to further analyses as described below.

### MR image acquisition, preprocessing, and analyses

Structural and functional brain imaging data were acquired with a Philips Achieva 3.0T MRI scanner (Philips Healthcare, The Netherlands) using a 32-channel head coil. A high-resolution 3D Turbo Field Echo T1-weighted image was acquired (3DTFE, TR = 9.7ms, TE = 4.6ms, field of view = 256 x 256 x 192 mm^3^, 192 sagittal slices, 1 x 1 x 1 mm voxel size). Functional images were acquired during visuomotor force tracking (see behavioral paradigm) with an ascending gradient echo-planar imaging (EPI) pulse sequence for T2*-weighted images using multiband factor imaging (repetition time = 1500ms, echo time =33ms, multiband factor = 2, flip angle = 80°, transversal slices = 40, slice thickness = 3mm, inter-slice gap = 0.5mm, matrix size 112 x 110 x 40 slices, 2.14 x 2.18 x 3.0 mm^3^ voxel dimensions, field of view =240 x 240 x 139.5mm^3^, 705 dynamic scans, excluding 4 dummy scans at the beginning of the sequence, SPIR fat suppression).

The fMRI data were preprocessed and analyzed using SPM12 (version 7771, Wellcome Department of Imaging Neuroscience, London, UK, http://www.fil.ion.ucl.ac.uk/spm) implemented in MATLAB version 9.3.0.713579 2017b (Natick, Massachusetts: The MathWorks Inc). During preprocessing, functional BOLD time series of each participant were realigned to each session’s first image and thereafter to the across-session mean image in an iterative process using rigid body transformations optimized to minimize the residual sum of squares between functional volumes. Mean functional images were then co-registered to the high-resolution T1-weighted anatomical image using a rigid body transformation optimized to maximize the normalized mutual information between the two images. Co-registration parameters were then applied to the realigned BOLD time series. Based on the co-registered and segmented (gray matter, white matter, cerebrospinal fluid, bone, soft tissue, and background) individual structural images, an average subject-based template was created using DARTEL in SPM12 and registered to the MNI space. All functional and anatomical images were then normalized using the resulting template. Finally, spatial smoothing was applied to all functional images (8-mm isotropic full-width at half-maximum, FWHM, Gaussian kernel). Head motion in six dimensions (i.e., rotations and translations in 3 planes) was quantified for each subject as a result of the functional realignment preprocessing step. Maximum absolute translations and frame-wise displacement^56^ of ≥ 2 voxels were defined as the exclusion criteria for excessive head motion during fMRI data acquisition for participants’ individual imaging runs. After excluding these artefactual runs (N = 16 in total), a total N=86 runs (sham N=31, 0° phase lag N=27, 180° phase lag N=28) were entered in the analysis, resulting in a varying number of runs per individual contrast as detailed in the results. In the remaining sample, we observed [given as ± median absolute deviation] absolute displacement across volumes acquired of 0.57 [±0.36] mm for translation, 0.54 [±0.46]° for rotation, and 0.57 [±0.32] mm for frame-wise displacement. None of these motion parameters systematically varied with stimulation condition (one-way ANOVA, all p>.2, Table s19).

### Task-related BOLD activity analysis

The statistical analysis followed two sequential steps accounting for fixed and random effects. In the first step, a general linear model (GLM) was computed on individual participant level by multiple regression of the voxel-wise time series onto a composite model containing the regressors of interest (fixed-effects analysis, FFX). The GLM included responses to the three task conditions (i.e., LEFT, RIGHT, BILATERAL) for each stimulation session (i.e., 0°, 180°, sham). Performance precision and time were modeled as parametric modulators. All regressors consisted of boxcar functions representing the on- and offset of the blocks of interest [15 seconds with trial numbers 3-17] and were convolved with the canonical hemodynamic response function. The task-free period between task blocks (Figure 1b) of 11±2 seconds allowed the hemodynamic response to relax and was modeled as implicit baseline in the block design. As for the behavioral analysis, signal acquired during ramping up and down of tACS, i.e., the first two and the last trials (-3 seconds in total), was not included in the analysis of interest but was, together with the task cues (1 sec. before ramping up), defined as separate ‘regressors of no interest’ modeled as short blocks. In addition, head-movement parameters were also modeled as regressors of no interest to account for BOLD signal changes that correlated with head movements. To remove low-frequency drift, the time series was high-pass filtered at 128 seconds cycle cut-off. Serial correlations were estimated using an autoregressive (1^st^ order) plus white noise model with a restricted maximum likelihood (ReML) algorithm. Linear contrasts on individual level were built to identify brain regions with systematic BOLD signal change through the interaction of TASK (within session) and STIMULATION CONDITION (between sessions). Linear contrasts involving the TASK by STIMULATION CONDITION interaction were constructed using the TASK contrasts as [LEFT vs RIGHT], [LEFT vs BILATERAL], and [RIGHT vs BILATERAL] and the STIMULATION CONTRASTS as [tACS_0°_ vs. sham], [tACS_180°_ vs. sham], and [tACS_0°_ vs. tACS_180°_]. As the main behavioral effects were observed for the [LEFT vs. RIGHT] conditions, the corresponding fMRI results are reported in the main text. For completeness, the fMRI results related to the interaction contrasts involving the BILATERAL condition are reported in the supplemental material (Tables s4, s5). The individual statistical parametric maps (SPM[T]) generated through these linear contrasts were further spatially smoothed (Gaussian kernel 6 mm FWHM) and fed into a second-level random-effects analysis generating group-level random effects maps (RFX) using t-statistics to allow for inferences across participants (one-sample t-test).

To confirm task-based brain activations within the expected sensorimotor network independent of stimulation, we ran a full-factorial model contrasting the three task conditions pooled over stimulation conditions. This analysis showed expected activations in a large network including bilateral pre- and postcentral cortex, cerebellar regions (lobule 4-5,6, 8), superior parietal cortex, right supplementary motor area, right thalamus, and left occipital cortex (Table s20). Furthermore, we ran two analyses confirming no unspecific stimulation effects (i.e., independent of phase) irrespective of task conditions firstly by pooling over both verum stimulation conditions (Table s21) and secondly by separately contrasting tACS0° and tACS180° against sham (Table s22).

### Functional connectivity analysis

To investigate how the modulation of the phase lag through tACS affected the functional connectivity between the stimulation targets and other brain regions, we performed psychophysiological interaction (PPI) analyses^22^. Here, we tested whether the stimulation targets, i.e., the right and the left primary motor cortices, systematically altered their task-related functional connectivity with the rest of the brain under the three stimulation conditions. For each individual and for each session, the first eigenvariate was extracted from the physiological time series using singular value decomposition of the time series across the voxels included in a 6-mm radius sphere centered on the M1 coordinates. Specifically, coordinates of the stimulation target seed regions in MNI space were (-34, -28, 54 mm) for the left primary motor cortex and (36, -22, 50 mm) for the right primary motor cortex. These coordinates were defined based on the peak activations on group level resulting from the left-versus right-hand task contrast pooled over all stimulation conditions at p<.05_FWE_ (whole brain and masked (using AAL) within left/right pre-central gyrus, clusters shown in Figure 4a, Table s23). For each session, one psychological regressor representing the contrast between two task variants was generated. A physiological regressor representing the activity in the seed area was also extracted for each session, as described above. For each session, the psychophysiological regressors resulted from the interaction between the psychological and the physiological regressors. To build this interaction regressor, the underlying BOLD activity was first estimated by a parametric empirical Bayes formulation, combined with the psychological factor, and subsequently convolved with the hemodynamic response function^57^. As described above for the analysis of BOLD activation, the GLM contained the same ‘regressors of no interest’. The linear contrasts testing for the effect of session (stimulation condition) on the psychophysiological interaction at the individual level were then further spatially smoothed (6-mm FWHM Gaussian kernel) and entered a second-level random-effects analysis, accounting for inter-subject variance (one-sample t-test).

### Regression analysis

To further investigate brain-behavior relationships, we ran separate second-level regression analyses between the BOLD activation and functional connectivity maps and the effector-specific (RMSE) and effector-independent (SmpEn) behavioral effects of tACS_0°_ mu-frequency stimulation. For this purpose, stimulation condition differences were computed for each hand separately for the two behavioral outcome parameters (i.e., ΔRMSE = RMSE_0°_ - RMSE_sham_, ΔSmpEn = SmpEn_0°_ - SmpEn_sham_) and used as covariates. Subsequently, individual activation- and connectivity-based contrast images testing for the TASK by STIMULATION contrast of LEFT versus RIGHT for tACS_0°_ versus sham were regressed against each of the behavioral change covariates, i.e., ΔRMSE and ΔSmpEn. Following up on the previous analysis steps, we focused on the RM1 as the seed region for PPI-based contrasts.

Finally, in a separate set of regression analyses, we investigated the association between the electrical field strength in the primary motor stimulation targets on the tACS-induced BOLD signal and functional connectivity (PPI) changes with tACS_0°_ compared to sham.

As described above (section *Electrical field simulation*), we used the individual electrical field strengths extracted for 10mm spheres around peak coordinates of the stimulation target seed regions in MNI space for left primary (-34, -28, 54 mm) and right (36, -22, 50 mm) primary motor cortex and used them as covariates (LM1 eField, RM1 eField) in separate regression analysis. Individual contrasts for left- or right-hand related BOLD activation changes under tACS_0°_ compared to sham were regressed against the eField covariates in separate analyses. Based on the PPI analyses, individual contrasts testing for tACS_0°_ vs sham for left- versus right-hand were likewise regressed against the eField covariates in separate analyses, using either LM1 or RM1 as seed region.

### Statistical inference

A voxel-wise threshold of p_unc_<.001 (uncorrected for multiple comparisons, minimum extent of 10 voxels) was used to define and display clusters in each analysis described above (activity and connectivity). Statistical inferences were performed at a threshold of p < 0.05 after family-wise error (FWE) correction for multiple comparisons over small spherical volumes (small volume correction, SVC, approach^58,59^). Published data is still inconsistent regarding tACS effects on BOLD activation relative to a given set of electrodes and stimulation frequency^60,61^. It supports, however, the hypothesis that tACS is more likely to modulate task-related neural activity beyond the cortical areas covered by the electrodes ^62–64^. Therefore, we based our a priori assumption concerning regions of interest with relevant activation changes on knowledge about the sensorimotor network involved in controlling visuomotor isometric force tracking for uni- and bimanual tasks. These previous studies report an extended network including visuomotor cortical circuits, cortico-striatal circuits for action selection and prediction of grip force, and cortico-cerebellar circuits for feedback-guided motor output^30,65^. Based on these a priori selected regions of interest (ROI), spheres of 10-mm radius were centered around coordinates retrieved from literature (Table s24). An additional correction for multiple testing was performed using Bonferroni correction procedures for the number of regions of interest within each contrast (p_Bonf_ < 0.05). For completeness, all global maxima surviving the correction steps are reported even when falling outside the predefined ROIs.

Activation maps were overlaid on the group mean structural image, and anatomical descriptions of significant clusters were determined using the automated anatomical labeling atlas (AAL version 4 25/08/2015)^66^.

## Acknowledgments

The authors thank René Clerckx and Paul Meugens for skillful technical support, Elle Bokken and Melissa Nigro for support with data acquisition, and all volunteers for their motivation to participate in the many hours required for this study.

This work was supported by the Research Foundation Flanders (FWO) (G089818N, G039821N), the Excellence of Science grant (EOS, 30446199, MEMODYN), and the KU Leuven Research Fund (C16/15/070). ND was supported by an MSCA Global Postdoctoral Fellowship (No. 101068893-MemUnited).

## Data and Code Availability

Source data and code to reproduce analyses and figures presented in the main manuscript and supplemental material will be made publicly available through Open Science Framework upon publication.

## Supplementary Information

### Individual mu peak frequency distribution

**Figure s1.**
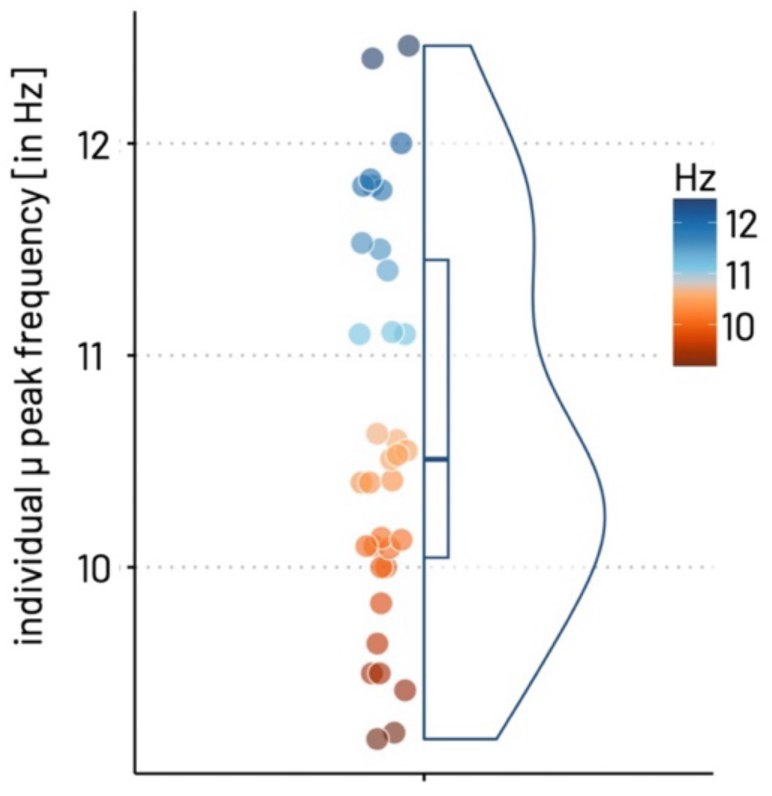
The individual mu peak frequency (iµPF) was estimated using a Gaussian curve fit^2^. Specifically, the highest local maximum within the mu frequency range ±1Hz (i.e., 7 – 14Hz) was estimated with 0.01 Hz resolution for both C3 and C4 electrodes supposed to capture signals from left and right primary motor cortices. Based on visual inspection, iµPF from the hemisphere that showed the most optimal fit was chosen as stimulation frequency (N = 35 from C3, N = 3 from C4). Points represent individual participants of the final sample (N=34). Distribution of individual mu peak frequency (in Hz) depicted (Mean = 10.65, SD = 0.92, Median = 10.51, MAD = 0.89, range: [9.19, 12.46]). Boxplots show the lowest/highest data points without outliers as lower/upper whiskers, lower/upper hinge reflect 25% / 75% of the data, and group median at the midline.

### Supplementary behavioral results

**Table s2.**
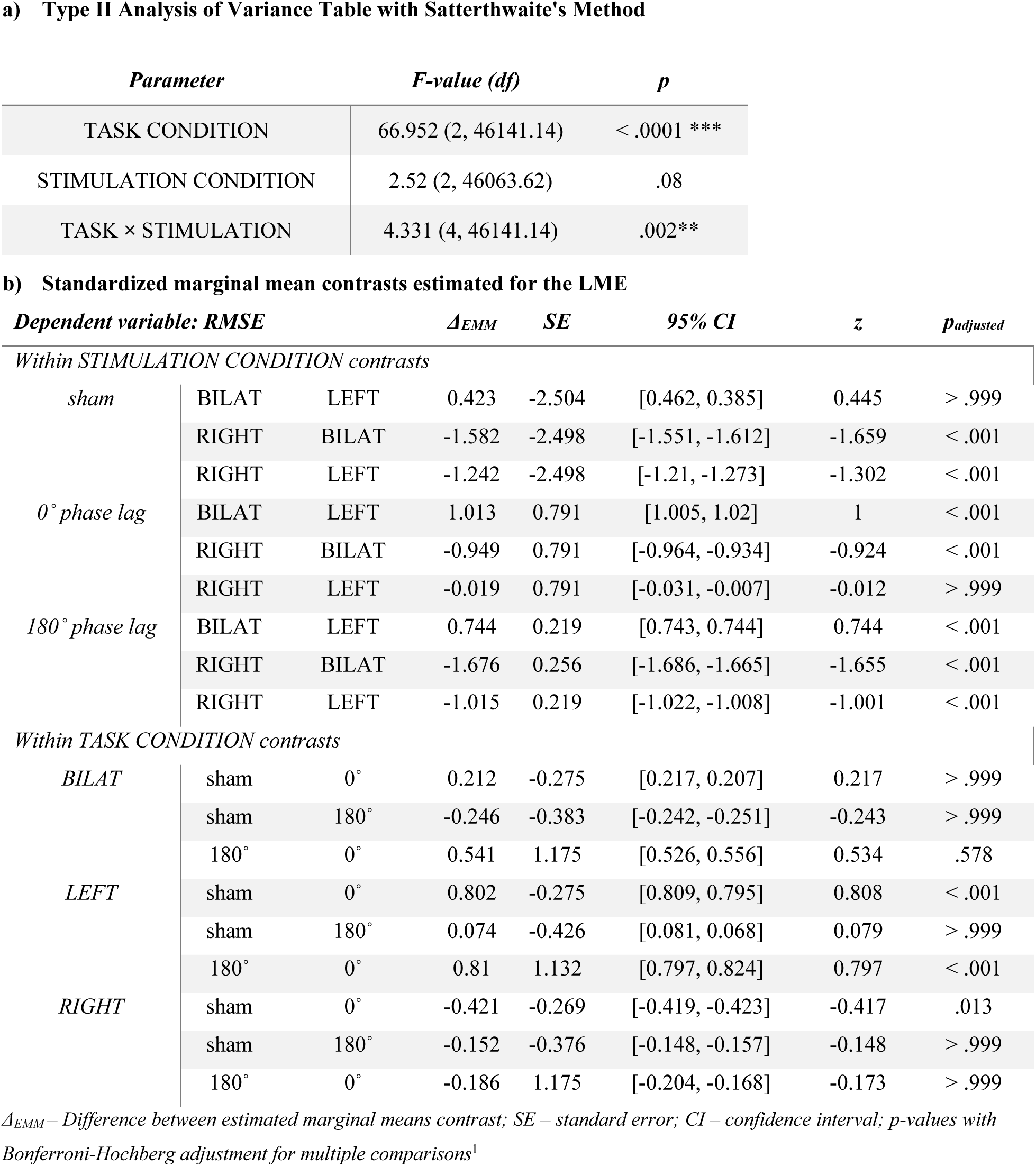
Linear mixed effects model (LME) results for RMSE.

**Table s3.** Linear mixed effects model (LME) results for SmpEn.

| *Parameter* | *F-value (df)* | *p* |
| --- | --- | --- |
| TASK CONDITION | 12.318 (2, 46140.95) | < .0001 *** |
| STIMULATION CONDITION | 206.081 (2, 46171.52) | < .0001 *** |
| TASK × STIMULATION | 2.677 (4, 46140.95) | 0.03* |

**b) Standardized marginal mean contrasts estimated for the LME**
| *Dependent variable: SmpEn* | | | $\Delta_{EMM}$ | *SE* | *95% CI* | *z* | *p<sub>adjusted</sub>* |
| --- | --- | --- | --- | --- | --- | --- | --- |
| *Within STIMULATION CONDITION contrasts* |  |  |  |  |  |  |  |
| *sham* | BILAT | LEFT | -1.847 | -2.511 | [-1.821, -1.873] | -1.869 | < .001 |
|  | RIGHT | BILAT | -1.005 | -2.505 | [-0.975, -1.035] | -0.983 | > .999 |
|  | RIGHT | LEFT | -1.57 | -2.505 | [-1.543, -1.597] | -1.578 | > .999 |
| *0° phase lag* | BILAT | LEFT | -1.418 | 0.781 | [-1.435, -1.401] | -1.409 | > .999 |
|  | RIGHT | BILAT | -0.547 | 0.781 | [-0.56, -0.534] | -0.552 | < .001 |
|  | RIGHT | LEFT | -0.684 | 0.781 | [-0.697, -0.67] | -0.686 | < .001 |
| *180° phase lag* | BILAT | LEFT | -1.708 | 0.209 | [-1.718, -1.697] | -1.698 | > .999 |
|  | RIGHT | BILAT | -0.481 | 0.247 | [-0.486, -0.475] | -0.478 | < .001 |
|  | RIGHT | LEFT | -0.906 | 0.209 | [-0.913, -0.899] | -0.901 | > .999 |
| *Within TASK CONDITION contrasts* |  |  |  |  |  |  |  |
| *BILAT* | sham | 0° | 1.215 | -0.274 | [1.225, 1.206] | 1.234 | < .001 |
|  | sham | 180° | -0.353 | -0.376 | [-0.349, -0.356] | -0.339 | < .001 |
|  | 180° | 0° | 0.286 | 1.176 | [0.271, 0.301] | 0.256 | < .001 |
| *LEFT* | sham | 0° | 1.644 | -0.274 | [1.655, 1.633] | 1.665 | < .001 |
|  | sham | 180° | -0.213 | -0.419 | [-0.208, -0.218] | -0.198 | < .001 |
|  | 180° | 0° | 0.576 | 1.133 | [0.562, 0.588] | 0.539 | < .001 |
| *RIGHT* | sham | 0° | 0.758 | -0.267 | [0.765, 0.751] | 0.774 | < .001 |
|  | sham | 180° | -0.877 | -0.369 | [-0.876, -0.878] | -0.867 | > .999 |
|  | 180° | 0° | 0.353 | 1.176 | [0.338, 0.367] | 0.321 | < .001 |

$\Delta_{EMM}$ – Difference between estimated marginal means contrast; *SE* – standard error; *CI* – confidence interval; *p*-values with Bonferroni-Hochberg adjustment for multiple comparisons<sup>1</sup>

**Figure s11.**
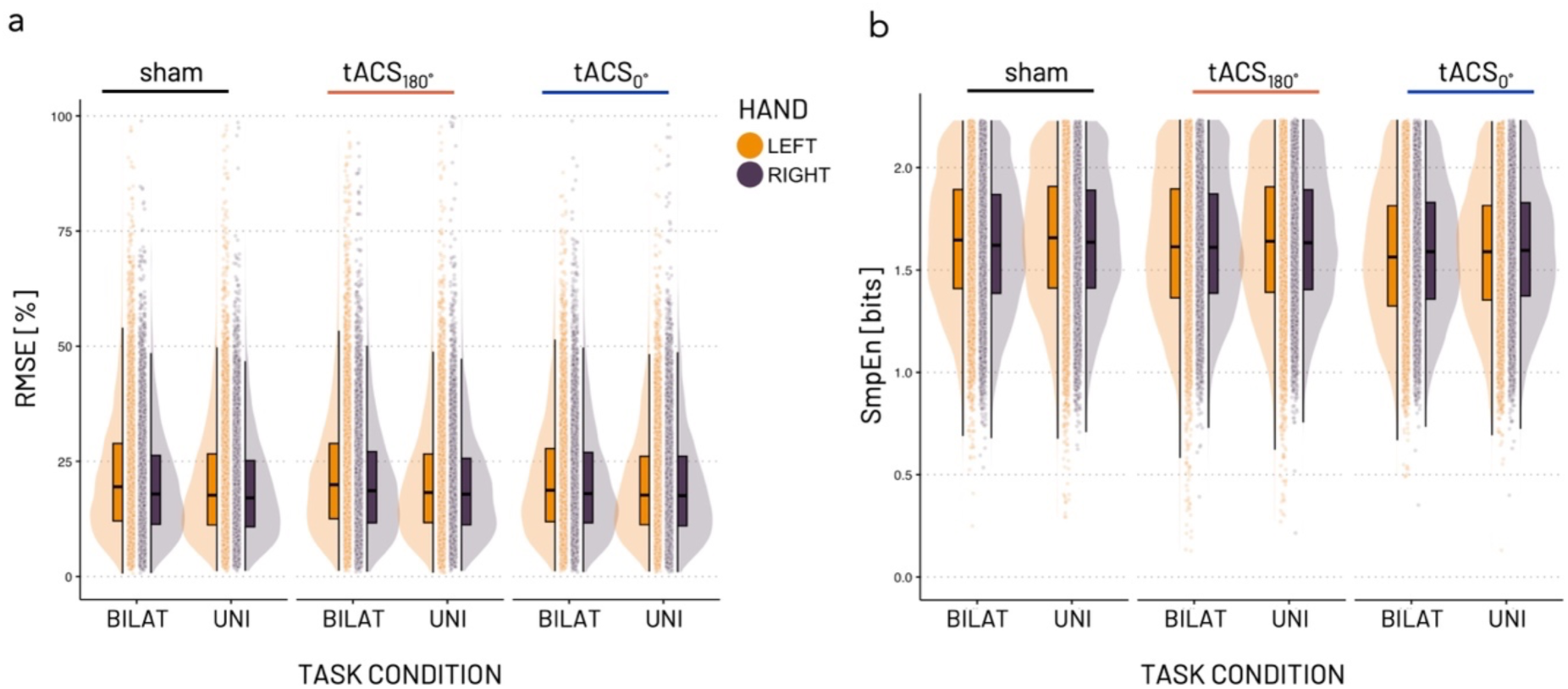
Comparison of unilateral versus bilateral behavioral outcomes for the left (orange) and right (purple) hand. a) Root Mean Square Error (RMSE) of force adjustment is generally higher, i.e., more erroneous in the non-dominant left hand, independent of uni- or bilateral task condition. While the bilateral RMSE is overall slightly higher than the unilateral RMSE for the left and right hand, the distributions remain the same. b) Overall, Sample Entropy (SmpEn) of force adjustment is slightly lower in the bilateral task condition. Points depict individual trials. Boxplots show the lowest/highest data points without outliers as lower/upper whiskers, lower/upper hinge reflect 25% / 75% of the data, and group median at the midline. Based on this data, left- and right-hand data were averaged for bilateral task conditions for all statistics reported in the manuscript to allow for comparison with the MRI data.

### Supplementary BOLD activation results

#### a) Contrasts involving bimanual task condition

**Table s4.** Bimanual versus left-hand action.

| *Anatomical region* | *Cluster extension (k)* | *Peak MNI coordinates (x, y, z mm)* | *Cluster Z score* | *Cluster $p_{FWE-SVC}$* |
| --- | --- | --- | --- | --- |
| **a) [L-B]<sub>0°tACS</sub>>[L-B]<sub>sham</sub>** |  |  |  |  |
| no clusters |  |  |  |  |
| **b) [L-B]<sub>sham</sub>>[L-B]<sub>0°tACS</sub>** |  |  |  |  |
| no clusters |  |  |  |  |
| **c) [L-B]<sub>180°tACS</sub>>[L-B]<sub>sham</sub>** |  |  |  |  |
| no clusters |  |  |  |  |
| **d) [L-B]<sub>sham</sub>>[L-B]<sub>180°tACS</sub>** |  |  |  |  |
| Thalamus, left | 156 | -14, -8, 10 | 3.72 | .004 |
|  |  | -18, -18, 10 | 3.54 | .007 |
| Calcarine cortex | 17 | 0, -94, -2 | 3.59 | .006 |
| Cerebellum, lobule IV/V, left | 161 | -30, -42, -24 | 3.75 | .003* |
|  |  | -38, -44, -22 | 3.53 | .007 |
| Cerebellar Vermis (IV-V) | 58 | -2, -64, -6 | 3.85 | .002* |
| Thalamus VL, left<br>Caudate, left | 69 | -14, -8, 12 | 3.67 | .004 |
|  |  | -14, -4, 18 | 3.66 | .005 |
|  |  | -12, 0, 18 | 3.55 | .006 |
| Medial Frontal Gyrus/anterior prefrontal cortex, left | 40 | -36, 38, 22 | 3.70 | .004 |
| Cerebellum III-V, right | 24 | 12, -44, -22 | 3.56 | .006 |
| Ant Ins | 56 | -28, 18, 12 | 3.60 | .006 |
| Postcentral gyrus, left | 11 | -18, -34, 68 | 3.35 | .012 |
| **e) [L-B]<sub>180°tACS</sub>>[L-B]<sub>0°tACS</sub>** |  |  |  |  |
| no clusters |  |  |  |  |
| **f) [L-B]<sub>0°tACS</sub>>[L-B]<sub>180°tACS</sub>** |  |  |  |  |
| Middle cingulate cortex, left | 141 | -6, -8, 30 | 4.96 | .00003** |
|  |  | 10, -6, 32 | 4.38 | .0003** |
| Caudate, left | 14 | -24, -12, 22 | 3.57 | .006* |
| Inferior Frontal Gyrus Pars Triangularis, left | 37 | -44, 34, 0 | 3.42 | .009 |
| Cerebellar Vermis (VI) | 60 | 0, -74, -16 | 3.52 | .006* |
| Middle Frontal<br>Gyrus/Frontal Pole left | 86 | -22, 48, 20 | 3.51 | .007* |

Asterisks indicate adjusted *p*-values after additional multiple comparison correction for clusters within each contrast at $p_{Bonf} < .01^{**}$ , $p_{Bonf} < .05^{*}$

**Table s5.**
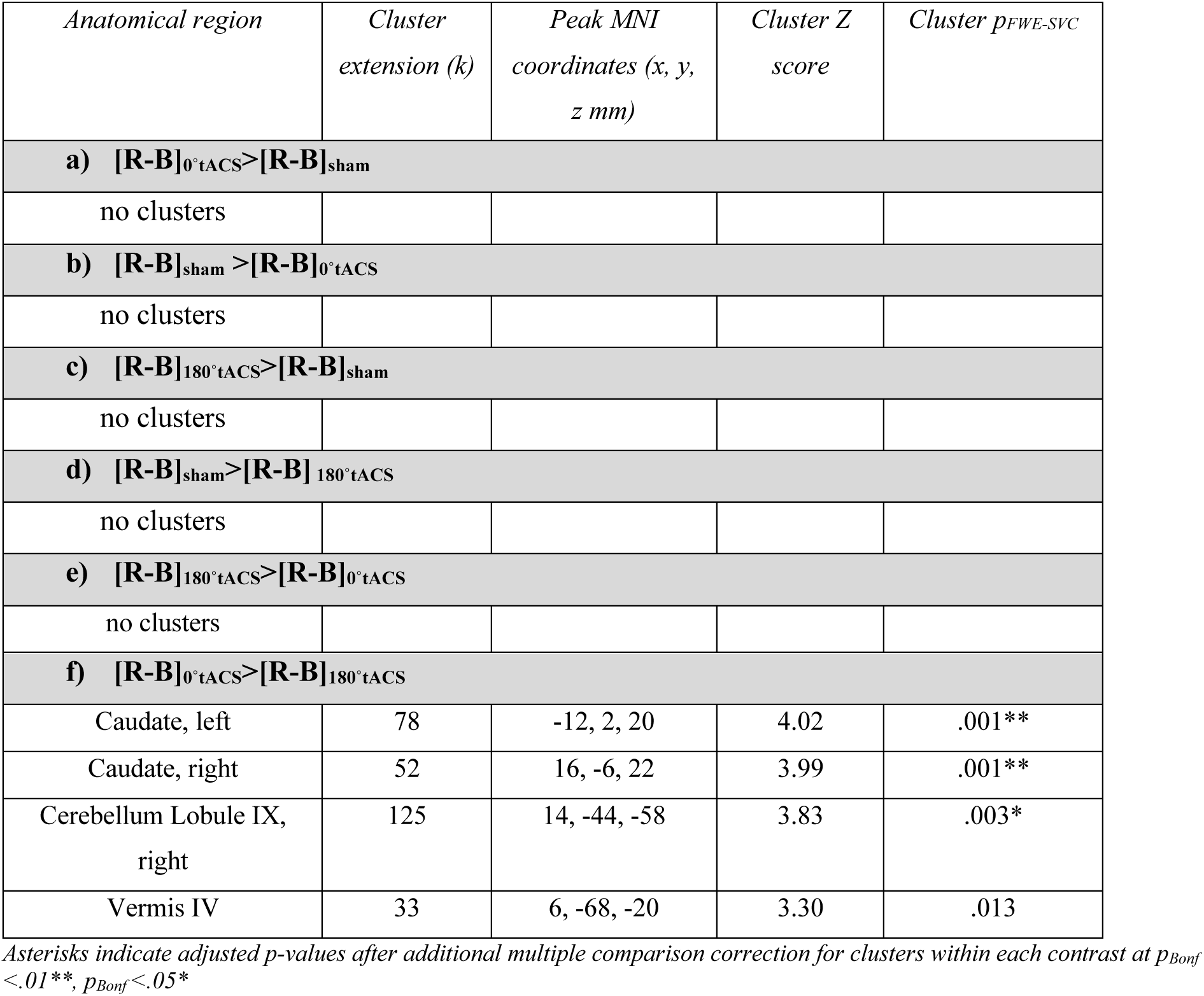
Bimanual versus right-hand action.

| *Anatomical region* | *Cluster extension (k)* | *Peak MNI coordinates (x, y, z mm)* | *Cluster Z score* | *Cluster $p_{FWE-SVC}$* |
| --- | --- | --- | --- | --- |
| **a) [R-B]<sub>0°tACS</sub> > [R-B]<sub>sham</sub>** |  |  |  |  |
| no clusters |  |  |  |  |
| **b) [R-B]<sub>sham</sub> > [R-B]<sub>0°tACS</sub>** |  |  |  |  |
| no clusters |  |  |  |  |
| **c) [R-B]<sub>180°tACS</sub> > [R-B]<sub>sham</sub>** |  |  |  |  |
| no clusters |  |  |  |  |
| **d) [R-B]<sub>sham</sub> > [R-B]<sub>180°tACS</sub>** |  |  |  |  |
| no clusters |  |  |  |  |
| **e) [R-B]<sub>180°tACS</sub> > [R-B]<sub>0°tACS</sub>** |  |  |  |  |
| no clusters |  |  |  |  |
| **f) [R-B]<sub>0°tACS</sub> > [R-B]<sub>180°tACS</sub>** |  |  |  |  |
| Caudate, left | 78 | -12, 2, 20 | 4.02 | .001** |
| Caudate, right | 52 | 16, -6, 22 | 3.99 | .001** |
| Cerebellum Lobule IX, right | 125 | 14, -44, -58 | 3.83 | .003* |
| Vermis IV | 33 | 6, -68, -20 | 3.30 | .013 |

Asterisks indicate adjusted *p*-values after additional multiple comparison correction for clusters within each contrast at $p_{Bonf} < .01^{**}$ , $p_{Bonf} < .05^{*}$

#### b) Confirmatory analysis without extreme values

**Table s6.** Results without extreme values.

| *Anatomical region* | *Cluster extension (k)* | *Peak MNI coordinates (x, y, z mm)* | *Cluster Z score* | *Cluster $p_{uncorrected}$* |
| --- | --- | --- | --- | --- |
| **a) tACS<sub>180°</sub> vs sham** |  |  |  |  |
| **[L-R]<sub>180°tACS</sub> > [L-R]<sub>sham</sub>** |  |  |  |  |
| no clusters |  |  |  |  |
| **[L-R]<sub>sham</sub> > [L-R]<sub>180°tACS</sub>** |  |  |  |  |
| Hippocampus, left | 246 | -26, -38, -6 | 4.54 | .000003* |
| Putamen, left | 795 | -22, 12, -6 | 4.10 | .00002 |
|  |  | -20, -4, 12 | 3.90 | .00005 |
| Hippocampus, right | 85 | 24, -36, -2 | 3.82 | .00007 |
| Putamen, right | 170 | 30, 8, -8 | 3.71 | .0001 |
| Cerebellar Vermis [VIII],<br>left | 64 | -2, -58, -28 | 3.50 | .0002 |

\* Whole-brain FWE correction at $p < .05$ .

#### c) Control analysis to test task-specific activation

The main effect of task condition (LEFT vs. RIGHT vs. BILATERAL) irrespective of the stimulation condition highlighted a large network including bilateral pre- and postcentral cortex, cerebellar regions (lobule 4-5,6, 8), superior parietal cortex, right supplementary motor area, right thalamus, and left occipital cortex.

**Table s20.** Main effect of TASK (LEFT vs RIGHT vs BIMANUAL)

| *Anatomical region* | *Cluster extension (k)* | *Peak MNI coordinates (x, y, z mm)* | *Peak F value* | *Peak Z score* | *p<sub>FWE-corr</sub>* |
| --- | --- | --- | --- | --- | --- |
| Precentral gyrus, right<br>Cerebellum lobule IV-VI<br>right<br>Superior Parietal cortex, right | 7758 | 34, -20, 50 | 76.38 | Inf | .0001 |
|  |  | 14, -54, -22 | 37.81 | 7.75 | .0001 |
|  |  | 26, -50, 70 | 33.82 | 7.35 | .0001 |
| Post- and Precentral, left | 1237 | -36, -24, 50 | 61.14 | Inf | .0001 |
| Cerebellum lobule IV-VI,<br>VIII, left<br>Lingual, left | 2277 | -14, -54, -20 | 53.59 | Inf | .0001 |
|  |  | -20, -58, -50 | 45.71 | Inf | .0001 |
|  |  | -12, -84, -6 | 17.03 | 5.17 | .002 |
| Superior Occipital cortex, left | 371 | -16, -90, 14 | 22.26 | 5.96 | .0001 |
| Supplementary Motor Area,<br>right | 74 | 10, -12, 52 | 18.80 | 5.45 | .0004 |
| Cerebellum lobule VIII, right | 80 | 16, -64, -50 | 17.57 | 5.26 | .001 |
| Thalamus, right | 62 | 20, -24, 12 | 16.21 | 5.03 | .003 |
| Superior Parietal, left<br>Precuneus, left | 41 | -22, -50, 62 | 13.90 | 5.62 | .019 |
|  |  | -20, -52, 70 | 13.78 | 5.60 | .021 |

Results of full-factorial model reported with whole-brain FWE correction at $p < .05$ and cluster extent threshold of 10 voxels.

#### d) Control analysis testing for unspecific effect of real stimulation

To test for an unspecific main effect of real tACS on MR signal, we ran a model comparing verum tACS (i.e., pooled over tACS_0°_ and tACS_180°_) against sham, irrespective of task condition. This analysis revealed an increased BOLD signal for verum stimulation compared to sham in medial and superior occipital, superior parietal, pre-and postcentral regions, and an increased BOLD signal in left Cuneus under sham compared to verum stimulation.

**Table s21.**
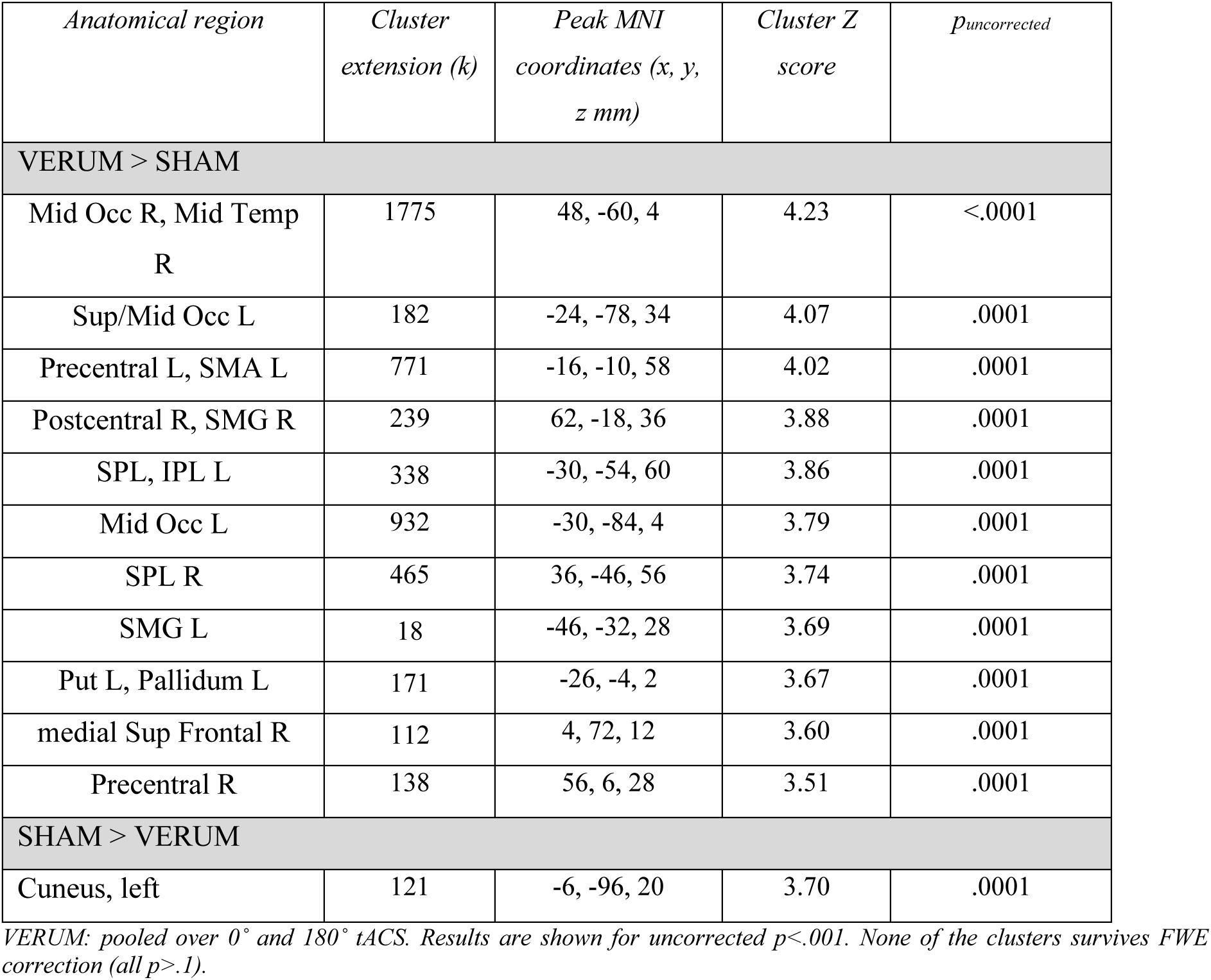
Effect of VERUM STIM vs. SHAM.

**Table s22.**
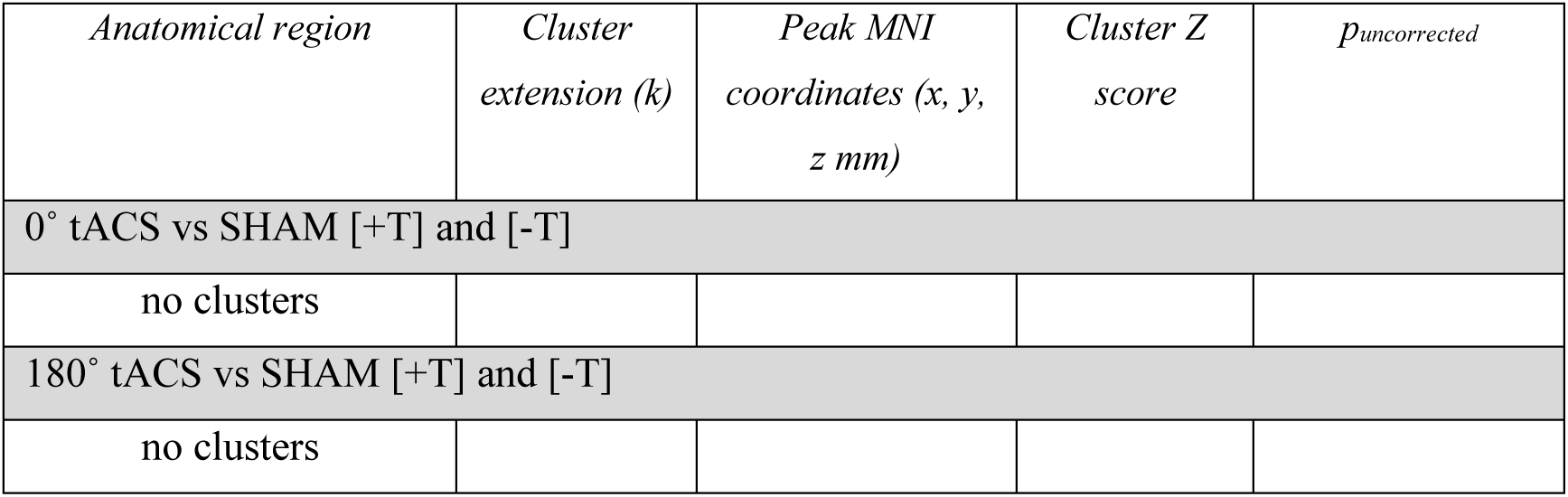
Pairwise test - effect of VERUM STIM vs. SHAM.

### Supplementary Regression results

#### a) Modelling behavioral outcomes as covariates

**Figure s7.**
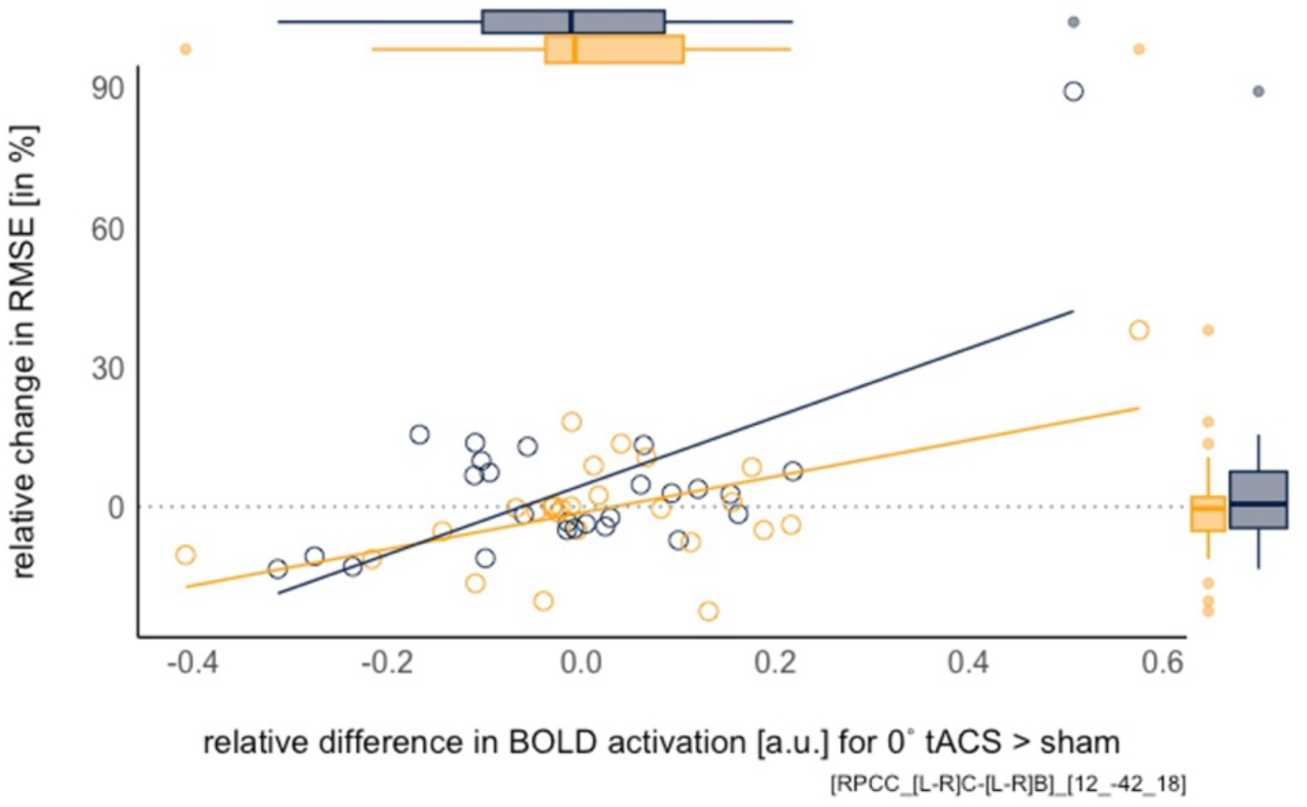
Regression results with right-hand RMSE as covariate. Association between right-hand RMSE increase and right PPC activation increase under tACS_0°_ compared to sham. Right hand is depicted in grey, left hand in yellow (note: left hand not significant in this contrast and added here for comparison only).

#### b) Modelling electrical field strength as covariate

**Figure s8.**
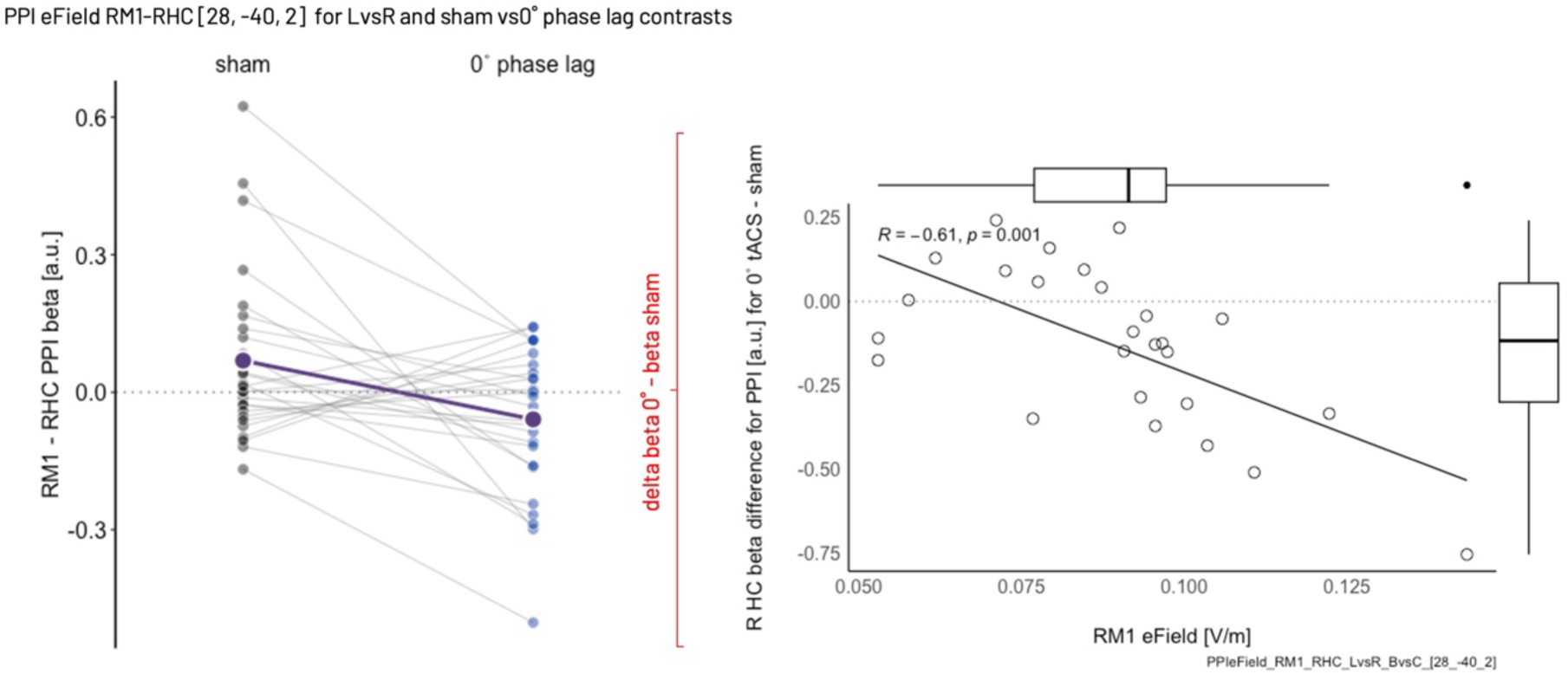
Regression results for RM1 eField. Association between higher eField strength in RM1 and more pronounced reduction in functional RM1- RHC connectivity **(negative difference of beta values tACS_0°_ - beta values sham) for left- versus right-hand contrast)**. **Left:** On average, individual functional connectivity between right M1 and right HC was reduced under 0° phase lag (blue) compared to sham condition (black/grey), leading to a negative difference (delta 0°- sham) as depicted on the y-axis of the scatter plot of Figure 5d (**right**). Purple points and slope depict group means.

**Table s9.** Regression results for LM1 eField. Associations between functional connectivity (PPI) of LM1 seed with the electrical field strength in the left M1

| *PPI (LM1 seed) – LM1 eField as covar – (+T) (LvsR sham vs 0°) [df:1, 24]* |  |  |  |  |
| --- | --- | --- | --- | --- |
| *Anatomical region* | *Cluster extension (k)* | *Peak MNI coordinates (x, y, z mm)* | *Z score* | *Cluster p<sub>FWE-SVC</sub>* |
| Hippocampus, right | 13 | 22, -38, 4 | 3.83 | .003* |
|  |  | 26, -38, 2 | 3.61 | .006* |
| Caudate, right | 49 | 22, 26, -4 | 3.92 | .002* |
|  |  | 20, 34, 10 | 3.29 | .015 |
| Caudate, left | 4 | -8, 24, 6 | 3.27 | .015 |
| Pallidum, right | 65 | 18, 4, 0 | 3.54 | .007* |

**Figure s10.**
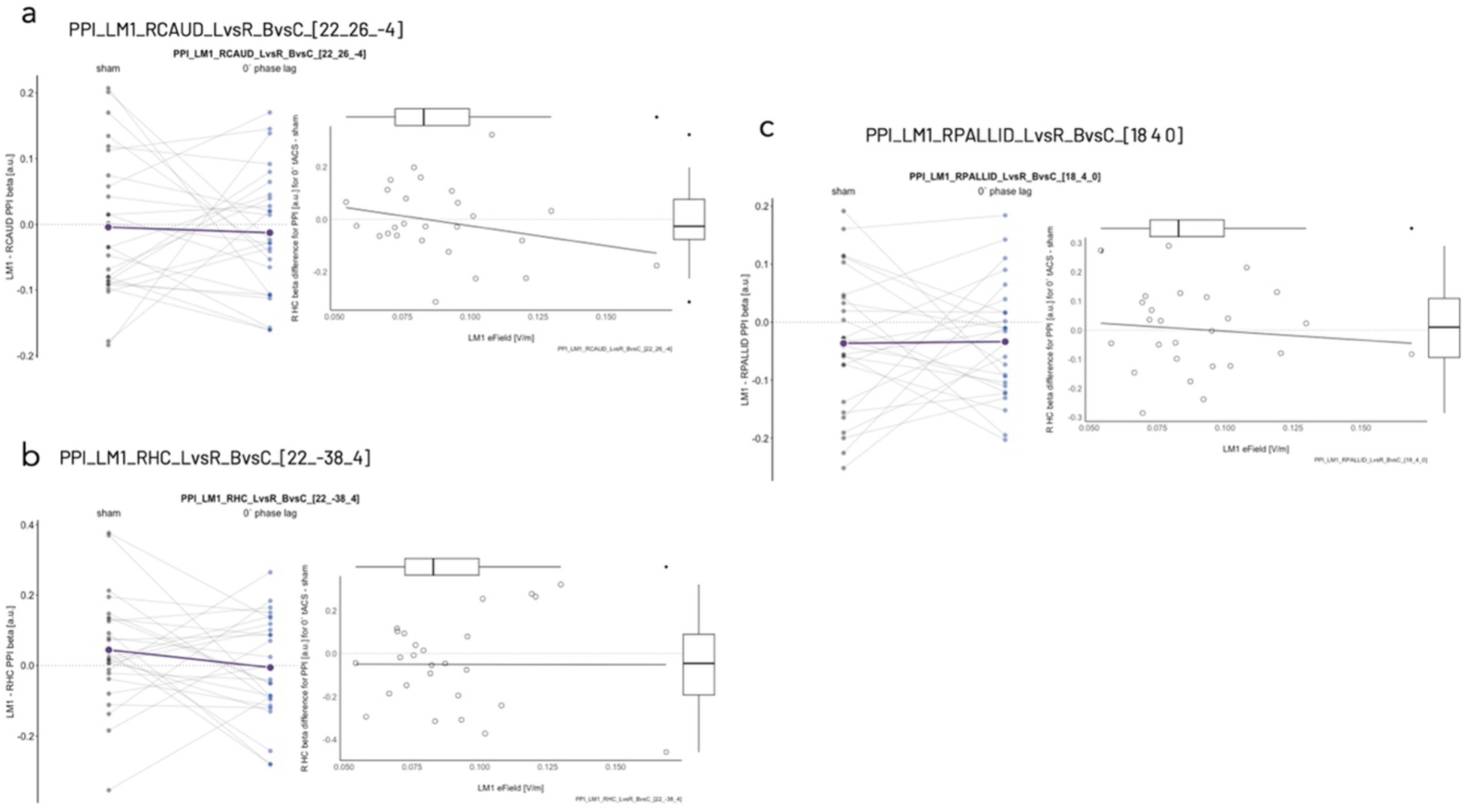
Regression results for LM1 eField. Associations between functional connectivity (PPI) of LM1 seed with the electrical field strength in the left M1 a) LM1-RCAUD, b) LM1-RHC, 3) LM1-RPALLIDUM

### Supplementary Results on Impedance values (tACS stimulation)

Bayesian general linear models (Gamma family with identity link) (estimated using MCMC sampling with 4 chains of 4000 iterations and a warmup of 2000) to predict Impedance with STIMULATION CONDITION (formula: Impedance ∼ STIMULATION CONDITION). Priors over parameters were all set as student_t (location = 0.00, scale = 1.00) distributions.

Following the Sequential Effect eXistence and sIgnificance Testing (SEXIT) framework, we report the median of the posterior distribution and its 95% CI (Highest Density Interval), along the probability of direction (pd), the probability of significance and the probability of being large. The thresholds beyond which the effect is considered as significant (i.e., non-negligible) and large are |0.05| and |0.30|. Convergence and stability of the Bayesian sampling has been assessed using R-hat, which should be below 1.01 (Vehtari et al., 2019), and Effective Sample Size (ESS), which should be greater than 1000 (Burkner, 2017). Constant monitoring of the stimulation electrode impedance revealed comparable impedance values across all stimulation conditions during the stimulation application. Expectedly, slightly higher impedance values were found in the sham condition due to higher maxima in the short ramp-in/out stimulation time of the sham condition. This effect was most prominent before the experiment outside the MRI (outside: median = 0.8kΩ, 95% CI [0.2, 11.8], inside: median = 11.2 kΩ, 95% CI [9.7, 13.7]), reflecting the expected impedance increase of 10kΩ due to resistors of MRI equipment (cables).

**Table s12.** Outside the MRI environment before the experiment.

|  | *Mean ±SD* | *Median [95%CI]* |
| --- | --- | --- |
| *sham* | 2.63±4.78 | 0.85 [0.2, 19.3] |
| *0° tACS* | 1.19±1.58 | 0.8 [0.2, 4.38] |
| *180° tACS* | 1.19±1.11 | 0.9 [0.3, 2.90] |

**b) Summary of posterior distributions**
| *Parameter* | *Median*<br>*[95% CI]*<br>*p (MAP)* | *pd*<br>*p (ROPE)*<br>*ps* | *[ROPE range]*<br>*% in ROPE*<br>*Equivalence (ROPE)* | *Rhat* | *ESS* | *BF* |
| --- | --- | --- | --- | --- | --- | --- |
| *(Intercept)* | 2.49<br>[ 2.00, 3.03]<br>< .001 | 100%<br>< .001<br>1.00 | [-0.31, 0.31]<br>0%<br>Rejected | 1.001 | 3380.00 | > 1000 |
| *0° tACS* | -1.26<br>[-1.90, -0.70]<br>.001 | 99.98%<br>.001<br>1.00 | [-0.31, 0.31]<br>0%<br>Rejected | 1.001 | 2889.00 | 340.12 |
| *180° tACS* | -1.26<br>[-1.88, -0.68]<br>< .001 | 100%<br>< .001<br>1.00 | [-0.31, 0.31]<br>0%<br>Rejected | 1.000 | 2813.00 | > 1000 |

MAP - maximum a posteriori probability estimate; pd- probability of direction; ESS -Effective Sample Size, CI - Highest Density Interval; ROPE - Region of Practical Equivalence; BF – Bayes Factor

**Table s13.**
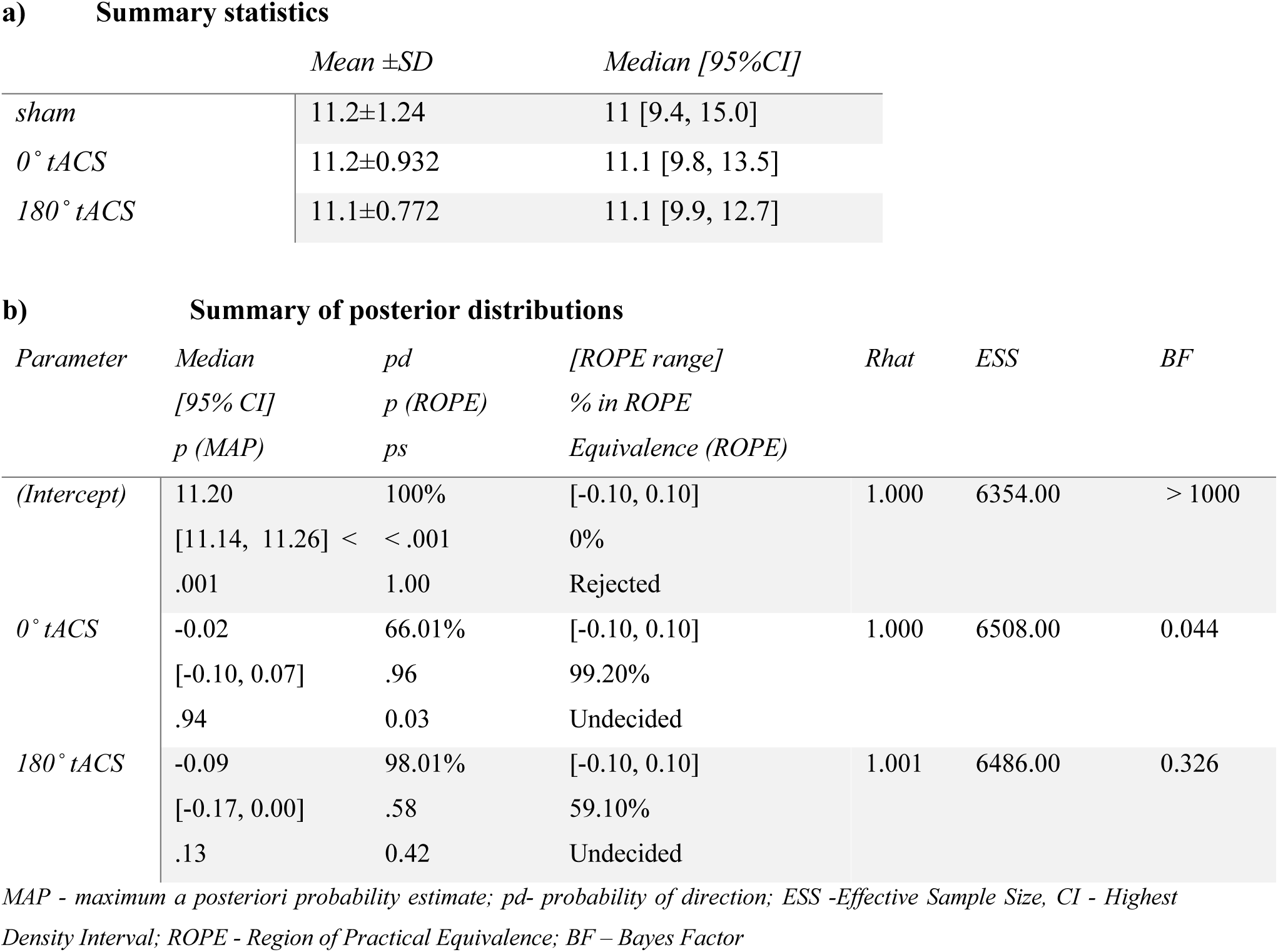
Inside the MRI environment during the experiment.

**Table s14.**
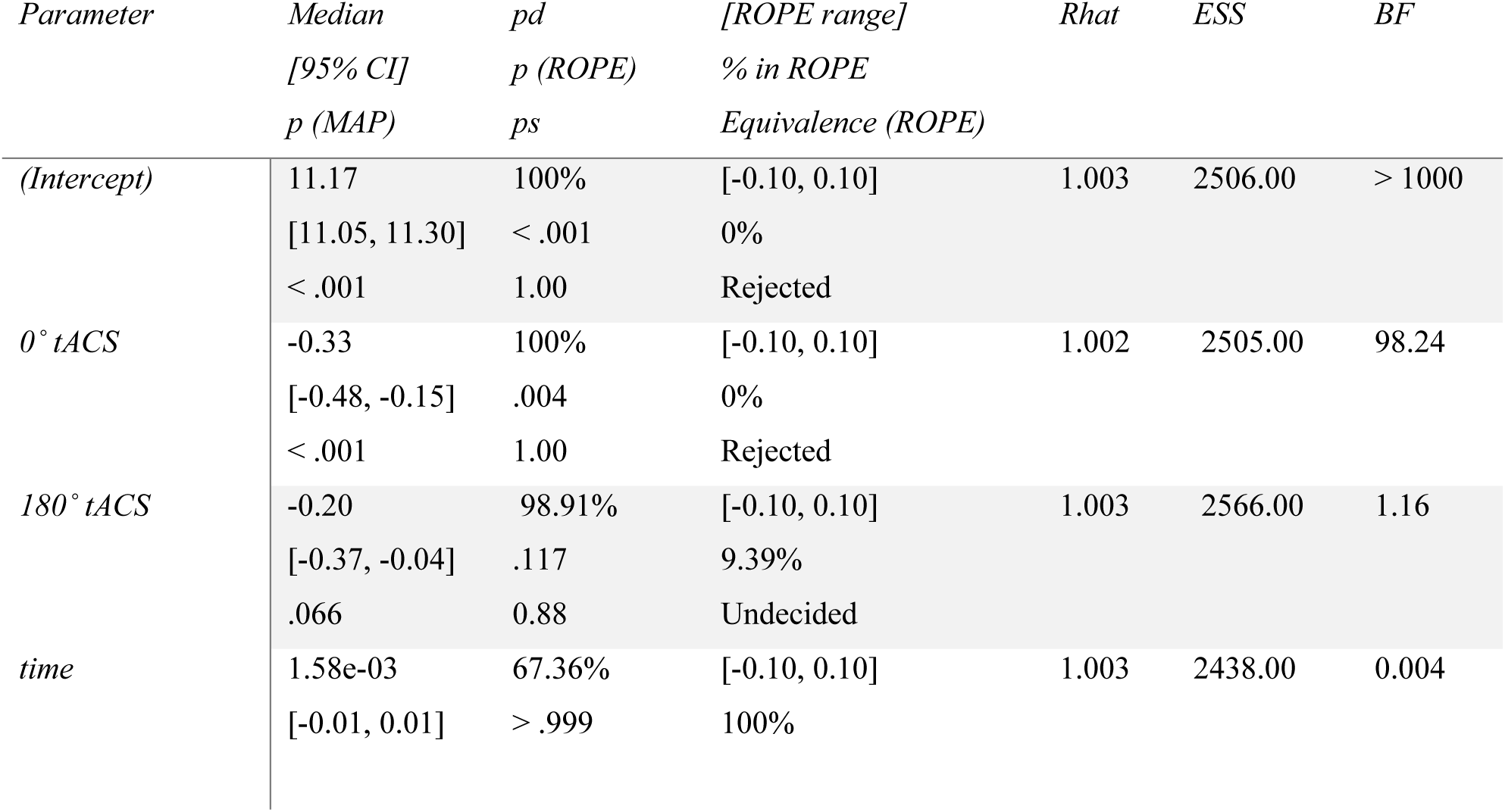
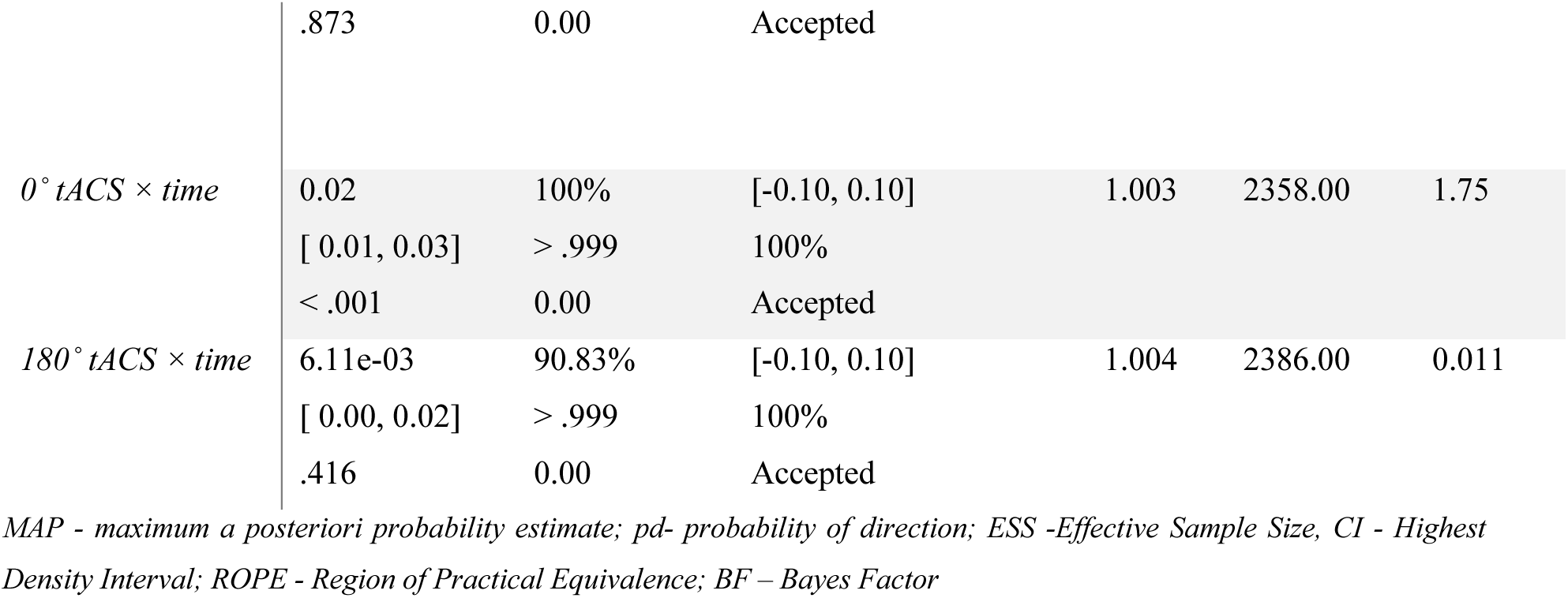
Inside the MRI environment during the experiment - effect of time Summary of posterior distributions.

### Supplementary information on neurosensory side effects and stimulation condition estimation

#### a) Subjective evaluation of neurosensory side effects

The level of the perceived strength of side effects was evaluated with a standardized questionnaire^3,4^ after each session. Bayesian probability tests were used to compare the hypothesis of equal distributions of relative frequencies of the estimations regarding the intensity (*None, Mild, Moderate, Considerable, Strong*) of perceived neurosensory side effects (Burning, Fatigue, Itching, Metallic Taste, Warmth, Pinching, Other) for each sub-category separately.

**Figure s15.**
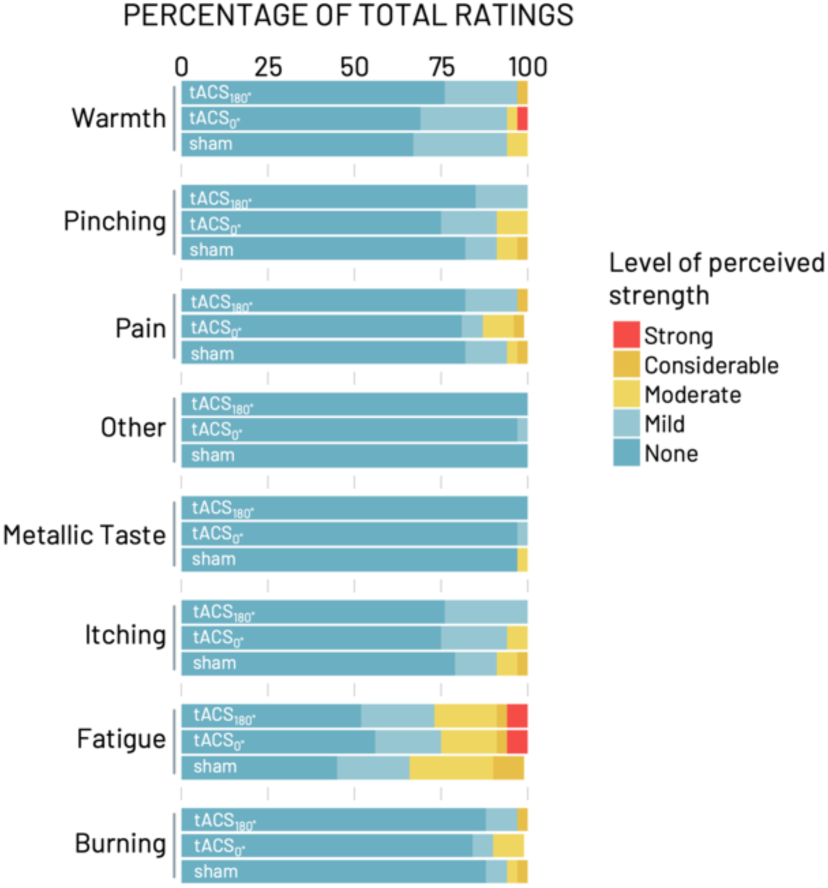
Level of the perceived strength of side effects. shown as the percentage of all submitted ratings for each category of side effect (y-axis) and stimulation condition. Color coding indicates the level of strength, with warmer colors representing more intense perception. Please note: Due to missing data (i.e., participants skipped categories when filling in the questionnaire), there are fewer overall ratings in the categories *Fatigue* for the sham and *Pain* and *Burning* for the tACS_0°_ condition.

For complexity reduction, probabilities were estimated for absent side effects (‘None’) and perceived side effects of any level of intensity (i.e., pooled over *Mild, Moderate, Considerable, Strong*). This analysis steps revealed that none of the stimulation conditions was more likely than the other two conditions to lead to an evaluation of 1) *None* or 2) *Any level of intensity* of neurosensory side effects.

**Table s16.**
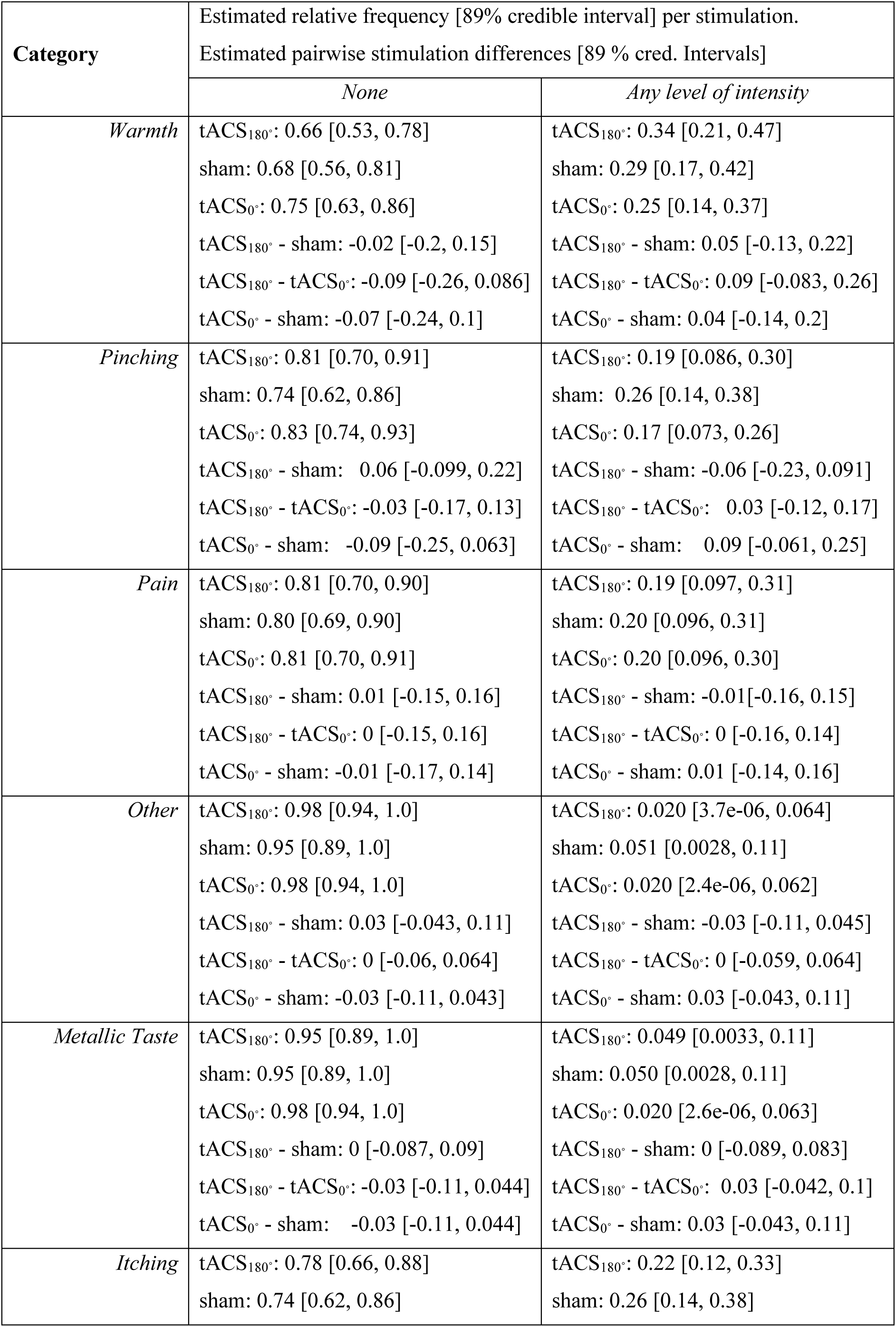
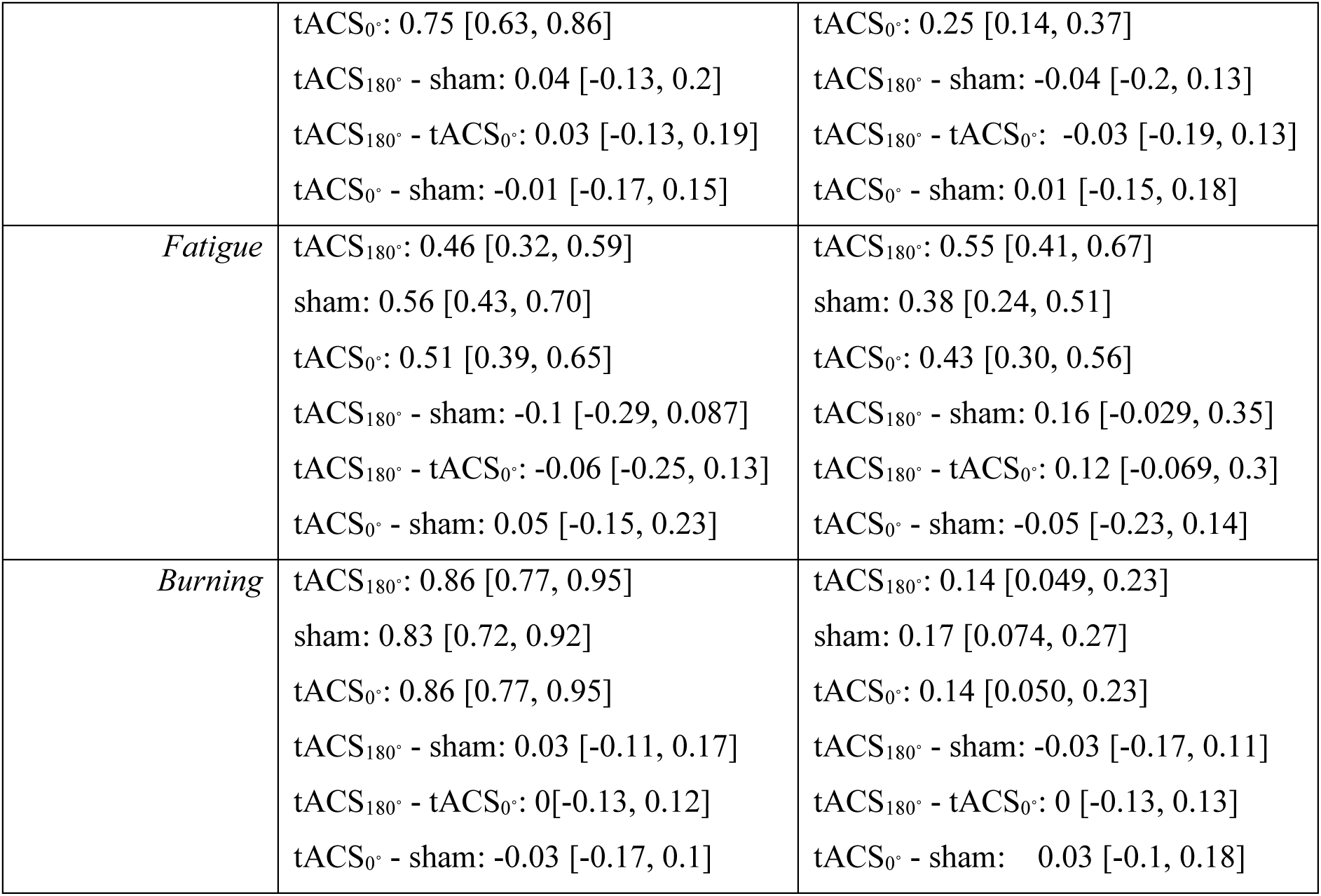
Results for Bayesian Probability Test Neurosensory Side Effects Estimation.

#### b) Post-hoc stimulation condition estimation

Bayesian probability tests comparing the hypothesis of equal distributions of relative frequencies of the estimations regarding the stimulation conditions verum (*It was real*), sham (*It was sham*), or no differentiation (*I don’t know*). Bayesian probability tests were run in JAGS, implemented for R^5,6^.

**Table s17.** Results Probability for Stimulation Condition Estimation.

| *Evaluation:* | *180° tACS vs sham* | *180° tACS vs 0° tACS* | *sham vs 0° tACS* |
| --- | --- | --- | --- |
| *I don't know* | pd = 0.96<br>p (MAP) = .22 | pd = 0.67<br>p (MAP) = .87 | pd = 0.91<br>p (MAP) = .38 |
| *It was real* | pd = 0.51<br>p (MAP) = 1.00 | pd = 0.96<br>p (MAP) = .20 | pd = 0.97<br>p (MAP) = .20 |
| *It was sham* | pd = 0.99*<br>p (MAP) = .07 | pd = 1.00*<br>p (MAP) = .01 | pd = 0.69<br>p (MAP) = .89 |

**Figure s18.**
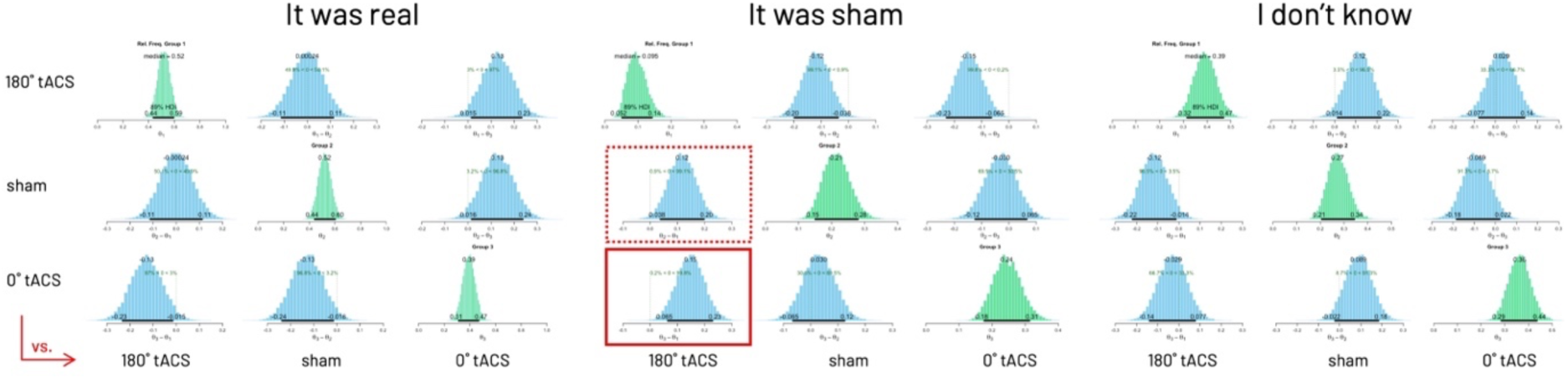
Based on the stimulation estimation after the last session, we found no evidence to support a difference in the participants’ evaluation of any of the stimulation conditions as being real stimulation. However, participants were slightly more likely to categorize sham (0.99, 89% HDI [0.038- 0.20]) and tACS_0°_ (1.0, 89% HDI [0.065-0.23]) as being sham than tACS_180°_ (outlined in red). Posterior distributions for pairwise comparisons of subjective estimations (*It was real*, *It was sham*, *I don’t know*) for the three stimulation conditions (180° tACS, 0° tACS, sham) are shown in blue, the diagonal is (comparison of each method with itself) shown green. Here, vertical dotted lines mark 0, positive posterior directions indicate a higher likelihood of condition shown on y-axis vs. condition on x-axis.

### Supplementary fMRI methods

**Table 19.**
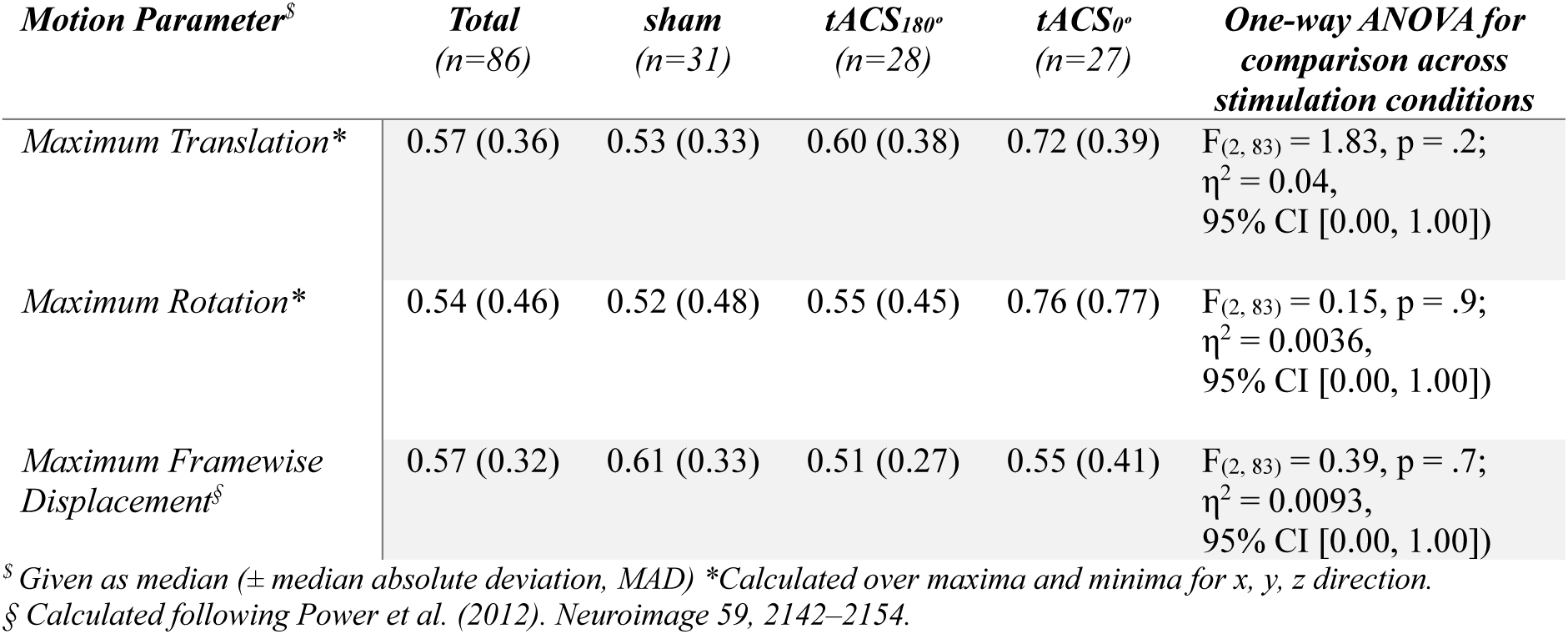
Motion parameters. Head motion in six dimensions (i.e., maximum rotations and linear translations across all three planes of movement) was compared with separate one-way ANOVAs for differences between STIMULATION CONDITIONS (sham, 0° phase lag, 180° phase lag tACS) for the final sample included in the analysis of BOLD time series:

**Table s23.** Definition of the seed coordinates for PPI and tACS targets. The coordinates of the stimulation target seed regions in MNI space were (-34, -28, 54 mm) for left primary motor cortex and (36, -22, 50 mm) for the right primary motor cortex. These coordinates were defined based on the peak activations on group level resulting from the left- versus right-hand task contrast pooled over all stimulation conditions at p<.05_FWE_ (whole brain and masked (using AAL) within left/right pre-central gyrus.

| *Anatomical region* | *Cluster*<br>*extension (k)* | *Peak MNI*<br>*coordinates (x, y,*<br>*z mm)* | *Cluster Z*<br>*score* | *P<sub>FWE</sub>* |
| --- | --- | --- | --- | --- |

| LEFT > RIGHT HAND [df: 1, 32] |  |  |  |  |
| --- | --- | --- | --- | --- |
| right precentral gyrus | 1223 | 36, -22, 50 | 5.58 | <.0001 |
|  |  | 20, -24, 76 | 5.09 | .003 |
|  |  | 24, -22, 68 | 5.07 | .004 |
| RIGHT > LEFT HAND [df: 1, 32] |  |  |  |  |
| left precentral gyrus | 465 | -34, -28, 54 | 5.54 | <.0001 |

Whole-brain FWE correction at $p < .05$ .

**Table s24.** Centroid coordinates used for small volume correction (SVC)

| *Anatomical region* | *Hemisphere* | *MNI coordinates*<br>(*x, y, z mm*) | *Literature*<br>*source* |
| --- | --- | --- | --- |
| *SFG/ DLPFC* | right | 23, 56, 30 | 7 |
| *Inferior Orbitofrontal Cortex* | right | 40, 26, -6* | 8 |
| *Medial Superior Frontal Gyrus* | right | 12, 46, 40* | 9 |
| *Caudate Nucleus* | left | -18, 18, 4* | 8 |
|  | right | 22, 26, 6 | 10 |
| *Putamen* | left | -28, 10, 2 | 11 |
|  | right | 24.2, 0.5, -6.1; 26, 10, 16* | 12,13 |
| *Hippocampus* | left | -26, -34, -6 | 14 |
|  | right | 22, -44, -4 | 15 |
| *Medial Temporal Gyrus* | left | -51, -51, -3 | 16 |
| *Precuneus* | right | 6, -58, 12* | 17 |
| *Inferior Parietal Lobule* | left | -56, -44, 44 | 18 |
| *Posterior Cingulate Cortex* | right | 6, -38, 18 | 19 |
| *Cerebellum* |  |  |  |
| *Vermis VIII* | left | -2, -64, -28 | 20 |
| *Crus I* | right | 38, -58, -25 | 21 |

\* Asterisks here indicate coordinates in Talairach space, which were converted to MNI space with https://bioimagesuiteweb.github.io/webapp/mni2tal.html

## Notes

### Competing Interest Statement

The authors have declared no competing interest.

## References

1. Rao, R. P. N. & Ballard, D. H. Predictive coding in the visual cortex: a functional interpretation of some extra-classical receptive-field effects. Nat Neurosci 2, 79–87 (1999).

2. Mehta, M. R. Neuronal Dynamics of Predictive Coding. Neuroscientist 7, 490–495 (2001).

3. Friston, K. & Kiebel, S. Predictive coding under the free-energy principle. Philosophical Transactions of the Royal Society B: Biological Sciences 364, 1211–1221 (2009).

4. Jakobs, O. et al. Effects of timing and movement uncertainty implicate the temporo-parietal junction in the prediction of forthcoming motor actions. NeuroImage 47, 667–677 (2009).

5. Singer, W. Neuronal Synchrony: A Versatile Code for the Definition of Relations? Neuron 24, 49–65 (1999).

6. Fries, P. A mechanism for cognitive dynamics: neuronal communication through neuronal coherence. Trends in Cognitive Sciences 9, 474–480 (2005).

7. Fries, P. Rhythms for Cognition: Communication through Coherence. Neuron 88, 220–235 (2015).

8. Stefanou, M.-I., Desideri, D., Belardinelli, P., Zrenner, C. & Ziemann, U. Phase Synchronicity of μ-Rhythm Determines Efficacy of Interhemispheric Communication Between Human Motor Cortices. J. Neurosci. 38, 10525–10534 (2018).

9. O’Reilly, C. & Elsabbagh, M. Intracranial recordings reveal ubiquitous in-phase and in- antiphase functional connectivity between homotopic brain regions in humans. J. Neurosci. Res. 99, 887–897 (2021).

10. Nikouline, V. V., Linkenkaer-Hansen, K., Huttunen, J. & Ilmoniemi, R. J. Interhemispheric phase synchrony and amplitude correlation of spontaneous beta oscillations in human subjects: a magnetoencephalographic study: Neuroreport 12, 2487–2491 (2001).

11. Swinnen, S. P. et al. Shared neural resources between left and right interlimb coordination skills: The neural substrate of abstract motor representations. NeuroImage 49, 2570–2580 (2010).

12. Bundy, D. T. & Leuthardt, E. C. The Cortical Physiology of Ipsilateral Limb Movements. Trends in Neurosciences 42, 825–839 (2019).

13. Hari, R. Action–perception connection and the cortical mu rhythm. in Progress in Brain Research vol. 159 253–260 (Elsevier, 2006).

14. Johnson, L. et al. Dose-dependent effects of transcranial alternating current stimulation on spike timing in awake nonhuman primates. Sci. Adv. 6, eaaz2747 (2020).

15. Riddle, J. & Frohlich, F. Targeting neural oscillations with transcranial alternating current stimulation. Brain Research 147491 (2021) doi:10.1016/j.brainres.2021.147491.

16. Richman, J. S. & Moorman, J. R. Physiological time-series analysis using approximate entropy and sample entropy. American Journal of Physiology-Heart and Circulatory Physiology 278, H2039–H2049 (2000).

17. Ficco, L. et al. Disentangling predictive processing in the brain: a meta-analytic study in favour of a predictive network. Sci Rep 11, 16258 (2021).

18. Siman-Tov, T. et al. Is there a prediction network? Meta-analytic evidence for a cortical- subcortical network likely subserving prediction. Neuroscience & Biobehavioral Reviews 105, 262–275 (2019).

19. Oldfield, R. C. The assessment and analysis of handedness: The Edinburgh inventory. Neuropsychologia 9, 97–113 (1971).

20. Flood, M. W. & Grimm, B. EntropyHub: An open-source toolkit for entropic time series analysis. PLoS ONE 16, e0259448 (2021).

21. Richman, J. S., Lake, D. E. & Moorman, J. R. Sample Entropy. in Methods in Enzymology vol. 384 172–184 (Academic Press, 2004).

22. Friston, K. J. et al. Psychophysiological and Modulatory Interactions in Neuroimaging. NeuroImage 6, 218–229 (1997).

23. Bastos, A. M., Lundqvist, M., Waite, A. S., Kopell, N. & Miller, E. K. Layer and rhythm specificity for predictive routing. Proc. Natl. Acad. Sci. U.S.A. 117, 31459–31469 (2020).

24. Ali, A., Ahmad, N., De Groot, E., Johannes Van Gerven, M. A. & Kietzmann, T. C. Predictive coding is a consequence of energy efficiency in recurrent neural networks. Patterns 3, 100639 (2022).

25. Gordon, N., Koenig-Robert, R., Tsuchiya, N., van Boxtel, J. J. & Hohwy, J. Neural markers of predictive coding under perceptual uncertainty revealed with Hierarchical Frequency Tagging. eLife 6, e22749 (2017).

26. Poldrack, R. A. & Packard, M. G. Competition among multiple memory systems: converging evidence from animal and human brain studies. Neuropsychologia 41, 245–251 (2003).

27. Haber, S. N. Corticostriatal circuitry. Dialogues in Clinical Neuroscience 18, 7–21 (2016).

28. Schiffer, A.-M. & Schubotz, R. I. Caudate Nucleus Signals for Breaches of Expectation in a Movement Observation Paradigm. Front. Hum. Neurosci. 5, (2011).

29. den Ouden, H. E. M., Daunizeau, J., Roiser, J., Friston, K. J. & Stephan, K. E. Striatal Prediction Error Modulates Cortical Coupling. Journal of Neuroscience 30, 3210–3219 (2010).

30. Prodoehl, J., Corcos, D. M. & Vaillancourt, D. E. Basal ganglia mechanisms underlying precision grip force control. Neuroscience & Biobehavioral Reviews 33, 900–908 (2009).

31. Sommer, S. & Pollmann, S. Putamen Activation Represents an Intrinsic Positive Prediction Error Signal for Visual Search in Repeated Configurations. TONIJ 10, 126–138 (2016).

32. Takada, M., Tokuno, H., Nambu, A. & Inase, M. Corticostriatal input zones from the supplementary motor area overlap those from the contra- rather than ipsilateral primary motor cortex. Brain Research 791, 335–340 (1998).

33. Lieu, C. A. & Subramanian, T. The interhemispheric connections of the striatum: Implications for Parkinson’s disease and drug-induced dyskinesias. Brain Research Bulletin 87, 1–9 (2012).

34. Ondobaka, S., de Lange, F. P., Wittmann, M., Frith, C. D. & Bekkering, H. Interplay Between Conceptual Expectations and Movement Predictions Underlies Action Understanding. Cerebral Cortex 25, 2566–2573 (2015).

35. Stachenfeld, K. L., Botvinick, M. M. & Gershman, S. J. The hippocampus as a predictive map. Nat Neurosci 20, 1643–1653 (2017).

36. Barron, H. C., Auksztulewicz, R. & Friston, K. Prediction and memory: A predictive coding account. Progress in Neurobiology 192, 101821 (2020).

37. Callaert, D. V. et al. Hemispheric asymmetries of motor versus nonmotor processes during (visuo)motor control. Hum. Brain Mapp. 32, 1311–1329 (2011).

38. Nee, D. E. Integrative frontal-parietal dynamics supporting cognitive control. eLife 10, e57244 (2021).

39. Du, J. et al. Functional connectivity of the orbitofrontal cortex, anterior cingulate cortex, and inferior frontal gyrus in humans. Cortex 123, 185–199 (2020).

40. Fine, J. M. & Hayden, B. Y. The whole prefrontal cortex is premotor cortex. Philosophical Transactions of the Royal Society B: Biological Sciences 377, 20200524 (2021).

41. Granek, J. A., Gorbet, D. J. & Sergio, L. E. Extensive video-game experience alters cortical networks for complex visuomotor transformations. Cortex 46, 1165–1177 (2010).

42. Barceló, F. A Predictive Processing Account of Card Sorting: Fast Proactive and Reactive Frontoparietal Cortical Dynamics during Inference and Learning of Perceptual Categories. Journal of Cognitive Neuroscience 33, 1636–1656 (2021).

43. Kasten, F. H., Dowsett, J. & Herrmann, C. S. Sustained Aftereffect of α-tACS Lasts Up to 70 min after Stimulation. Frontiers in Human Neuroscience 10, (2016).

44. Neuling, T., Rach, S. & Herrmann, C. Orchestrating neuronal networks: sustained after- effects of transcranial alternating current stimulation depend upon brain states. Frontiers in Human Neuroscience 7, (2013).

45. Hochberg, Y. A sharper Bonferroni procedure for multiple tests of significance. Biometrika 75, 800–802 (1988).

46. Bates, D., Mächler, M., Bolker, B. & Walker, S. Fitting Linear Mixed-Effects Models Using **lme4**. J. Stat. Soft. 67, (2015).

47. 47. Lüdecke, D., et al. Framework for Easy Statistical Modeling, Visualization, and Reporting. (2022).

48. 48. Mitra, P. & Bokil, H. Observed Brain Dynamics. (Oxford University Press, Oxford ; New York, 2008).

49. Gomez-Herrero, G. et al. Automatic Removal of Ocular Artifacts in the EEG without an EOG Reference Channel. in *Proceedings of the 7th Nordic Signal Processing Symposium - NORSIG* 2006 130–133 (IEEE, Rejkjavik, 2006). doi:10.1109/NORSIG.2006.275210.

50. Haegens, S., Cousijn, H., Wallis, G., Harrison, P. J. & Nobre, A. C. Inter- and intra- individual variability in alpha peak frequency. NeuroImage 92, 46–55 (2014).

51. Fertonani, A., Rosini, S., Cotelli, M., Rossini, P. M. & Miniussi, C. Naming facilitation induced by transcranial direct current stimulation. Behavioural Brain Research 208, 311– 318 (2010).

52. Fertonani, A., Ferrari, C. & Miniussi, C. What do you feel if I apply transcranial electric stimulation? Safety, sensations and secondary induced effects. Clinical Neurophysiology 126, 2181–2188 (2015).

53. Thielscher, A., Antunes, A. & Saturnino, G. B. Field modeling for transcranial magnetic stimulation: A useful tool to understand the physiological effects of TMS? in 2015 *37th Annual International Conference of the IEEE Engineering in Medicine and Biology Society (EMBC)* 222–225 (2015). doi:10.1109/EMBC.2015.7318340.

54. Puonti, O. et al. Accurate and robust whole-head segmentation from magnetic resonance images for individualized head modeling. NeuroImage 219, 117044 (2020).

55. Saturnino, G. B., Antunes, A. & Thielscher, A. On the importance of electrode parameters for shaping electric field patterns generated by tDCS. NeuroImage 120, 25–35 (2015).

56. Power, J. D., Barnes, K. A., Snyder, A. Z., Schlaggar, B. L. & Petersen, S. E. Spurious but systematic correlations in functional connectivity MRI networks arise from subject motion. Neuroimage 59, 2142–2154 (2012).

57. Gitelman, D. R., Penny, W. D., Ashburner, J. & Friston, K. J. Modeling regional and psychophysiologic interactions in fMRI: the importance of hemodynamic deconvolution. NeuroImage 19, 200–207 (2003).

58. Poldrack, R. A. Region of interest analysis for fMRI. Soc Cogn Affect Neurosci 2, 67–70 (2007).

59. Poldrack, R. A. et al. Guidelines for reporting an fMRI study. NeuroImage 40, 409–414 (2008).

60. Weinrich, C. A. et al. Modulation of Long-Range Connectivity Patterns via Frequency- Specific Stimulation of Human Cortex. Current Biology 27, 3061–3068.e3 (2017).

61. Moisa, M., Polania, R., Grueschow, M. & Ruff, C. C. Brain Network Mechanisms Underlying Motor Enhancement by Transcranial Entrainment of Gamma Oscillations. J. Neurosci. 36, 12053–12065 (2016).

62. Joundi, R. A., Jenkinson, N., Brittain, J.-S., Aziz, T. Z. & Brown, P. Driving Oscillatory Activity in the Human Cortex Enhances Motor Performance. Current Biology 22, 403–407 (2012).

63. Cabral-Calderin, Y., et al. Transcranial alternating current stimulation affects the BOLD signal in a frequency and task-dependent manner. Hum. Brain Mapp. 37, 94–121 (2016).

64. Vosskuhl, J., Huster, R. J. & Herrmann, C. S. BOLD signal effects of transcranial alternating current stimulation (tACS) in the alpha range: A concurrent tACS–fMRI study. NeuroImage 140, 118–125 (2016).

65. Floyer-Lea, A. & Matthews, P. M. Changing Brain Networks for Visuomotor Control With Increased Movement Automaticity. Journal of Neurophysiology 92, 2405–2412 (2004).

66. Tzourio-Mazoyer, N. et al. Automated Anatomical Labeling of Activations in SPM Using a Macroscopic Anatomical Parcellation of the MNI MRI Single-Subject Brain. NeuroImage 15, 273–289 (2002).

## Supplementary References

1. Hochberg, Y. A sharper Bonferroni procedure for multiple tests of significance. Biometrika 75, 800– 802 (1988).

2. Haegens, S., Cousijn, H., Wallis, G., Harrison, P. J. & Nobre, A. C. Inter- and intra-individual variability in alpha peak frequency. NeuroImage 92, 46–55 (2014).

3. Fertonani, A., Rosini, S., Cotelli, M., Rossini, P. M. & Miniussi, C. Naming facilitation induced by transcranial direct current stimulation. Behavioural Brain Research 208, 311–318 (2010).

4. Fertonani, A., Ferrari, C. & Miniussi, C. What do you feel if I apply transcranial electric stimulation? Safety, sensations and secondary induced effects. Clinical Neurophysiology 126, 2181–2188 (2015).

5. Plummer, M., Stukalov, A. & Denwood, M. rjags: Bayesian Graphical Models using MCMC. (2023).

6. Bååth, R. Bayesian First Aid. (2022).

7. Iannaccone, R. et al. Conflict monitoring and error processing: New insights from simultaneous EEG–fMRI. NeuroImage 105, 395–407 (2015).

8. Toni, I., Ramnani, N., Josephs, O., Ashburner, J. & Passingham, R. E. Learning Arbitrary Visuomotor Associations: Temporal Dynamic of Brain Activity. NeuroImage 14, 1048–1057 (2001).

9. de Jong, B. M. & Paans, A. M. J. Medial versus lateral prefrontal dissociation in movement selection and inhibitory control. Brain Research 1132, 139–147 (2007).

10. Zimmermann, M., Meulenbroek, R. G. J. & de Lange, F. P. Motor Planning Is Facilitated by Adopting an Action’s Goal Posture: An fMRI Study. Cerebral Cortex 22, 122–131 (2012).

11. Wenderoth, N., Debaere, F., Sunaert, S. & Swinnen, S. P. The role of anterior cingulate cortex and precuneus in the coordination of motor behaviour. European Journal of Neuroscience 22, 235–246 (2005).

12. Coombes, S. A., Corcos, D. M. & Vaillancourt, D. E. Spatiotemporal tuning of brain activity and force performance. NeuroImage 54, 2226–2236 (2011).

13. Lehéricy, S. et al. Motor control in basal ganglia circuits using fMRI and brain atlas approaches. Cerebral Cortex 16, 149–161 (2006).

14. Albouy, G. et al. Both the Hippocampus and Striatum Are Involved in Consolidation of Motor Sequence Memory. Neuron 58, 261–272 (2008).

15. Beudel, M., Leenders, K. L. & de Jong, B. M. Hippocampus activation related to ‘real-time’ processing of visuospatial change. Brain Research 1652, 204–211 (2016).

16. Kassuba, T. et al. The left fusiform gyrus hosts trisensory representations of manipulable objects. NeuroImage 56, 1566–1577 (2011).

17. Schulte, T. et al. fMRI evidence for individual differences in premotor modulation of extrastriatal visual–perceptual processing of redundant targets. NeuroImage 30, 973–982 (2006).

18. Callaert, D. V. et al. Hemispheric asymmetries of motor versus nonmotor processes during (visuo)motor control. Hum. Brain Mapp. 32, 1311–1329 (2011).

19. Ondobaka, S., de Lange, F. P., Wittmann, M., Frith, C. D. & Bekkering, H. Interplay Between Conceptual Expectations and Movement Predictions Underlies Action Understanding. Cerebral Cortex 25, 2566–2573 (2015).

20. Wenderoth, N., Toni, I., Bedeleem, S., Debaere, F. & Swinnen, S. P. Information processing in human parieto-frontal circuits during goal-directed bimanual movements. NeuroImage 31, 264–278 (2006).

21. Onuki, Y. Hippocampal–Cerebellar Interaction During Spatio-Temporal Prediction. Cereb. Cortex 9 (2015) doi:10.1093/cercor/bht221.

